# A chromosome-level reference genome of *Ensete glaucum* gives insight into diversity, chromosomal and repetitive sequence evolution in the Musaceae

**DOI:** 10.1101/2021.11.23.469474

**Authors:** Ziwei Wang, Mathieu Rouard, Manosh Kumar Biswas, Gaetan Droc, Dongli Cui, Nicolas Roux, Franc-Christophe Baurens, Xue-Jun Ge, Trude Schwarzacher, Pat (J.S.) Heslop-Harrison, Qing Liu

## Abstract

**Background:** *Ensete glaucum* (2*n* = 2*x* = 18) is a giant herbaceous monocotyledonous plant in the small Musaceae family along with banana (*Musa*). A high-quality reference genome sequence of *E. glaucum* offers a vital genomic resource for functional and evolutionary studies of *Ensete*, the Musaceae, and more widely in the Zingiberales.

**Findings:** Using a combination of Illumina and Oxford Nanopore Technologies (ONT) sequencing, genome-wide chromosome conformation capture (Hi-C), and RNA survey sequence, we report a high-quality assembly of the 481.5Mb genome with 9 pseudochromosomes and 36,836 genes (BUSCO 94.7%). A total of 55% of the genome is composed of repetitive sequences with LTR-retroelements (37%) and DNA transposons (7%) predominant. The 5S and 45S rDNA were each present at one locus, and the 5S rDNA had an exceptionally long monomer length of c.1,056 bp, contrasting with the c. 450 bp monomer at multiple loci in *Musa*. A tandemly repeated c. 134 bp satellite, 1.1% of the genome (with no similar sequence in *Musa*), was present around all nine centromeres, with a LINE retroelement also found at *Musa* centromeres. The assembly, including centromeric positions, enabled us to characterize in detail the chromosomal rearrangements occurring between the *x* = 9 species and *x* = 11 species of *Musa*. Only one chromosome has the same gene content as *M. acuminata* (ma). Three ma chromosomes represent part of only one *E. glaucum* (eg) chromosome, while the remaining seven ma chromosomes are fusions of parts of two, three, or four eg chromosomes, demonstrating complex and multiple evolutionary rearrangements in the change between *x* = 9 and *x* = 11.

**Conclusions:** The advance towards a Musaceae pangenome including *E. glaucum,* tolerant of extreme environments, makes a complete set of gene alleles available for crop breeding and understanding environmental responses. The chromosome-scale genome assembly show how chromosome number evolves, and features of the rapid evolution of repetitive sequences.

## Background

The genus *Ensete* Bruce ex Horaninow includes 10 species of giant, herbaceous monocotyledonous plants in the family Musaceae, native to tropical Africa and Asia [1]. Among them, the African species *E. ventricosum* (Welw.) Cheesman is an important food crop for more than 20 million people in Ethiopia [2]. Its sister genus *Musa,* grown throughout the tropics for food and fibre, includes diploid species, triploids and hybrids of *M. acuminata* and *M. balbisiana,* with banana cultivars. Sister to the grasses (Poales) and palms (Arecales) in monocots, both *Ensete* and *Musa,* along with a third genus *Musella,* belong to Musaceae in the order Zingiberales (gingers and bananas) [3]. Following rapid diversification of the Zingiberales at the Cretaceous/ Tertiary boundary (>65 Mya) the crown node age of the Musaceae family is soon after, with the *Musa* genus diverging from *Ensete* and *Musella* about 40 Mya [4, 5].

*Ensete glaucum* (Roxb.) Cheesman (NCBI:txid482298), like other species in *Ensete*, is monocarpic with a dilated and characteristically glaucous basal pseudostem, with a small number of large seeds (10 mm in diameter) in elongated, banana-like fruits, borne in hands with a terminal flower (Fig. 1A-E), and is diploid with 2*n* = 2*x* = 18 chromosomes [1,6–10]. *E. glaucum* is widely distributed in Asia and has records from Burma, China, India, Indonesia, India, Indonesia, Laos, Myanmar, Vietnam, Philippine, Papua New Guinea, Thailand, Solomon Islands [11].

**Figure 1.**
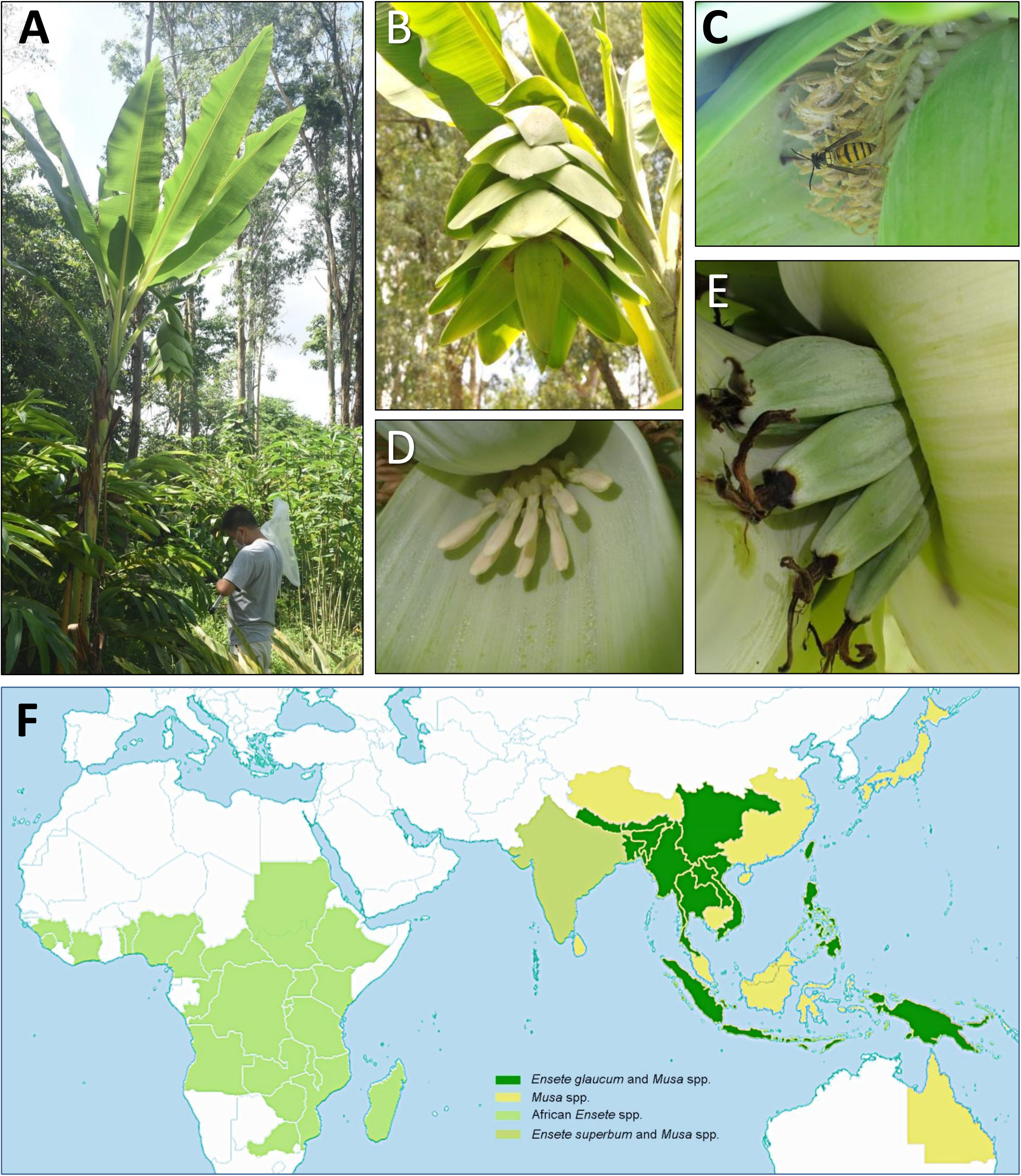
*Ensete glaucum* plant morphology and distribution map. (A) *E. glaucum* in South China Botanical Garden, Chinese Academy of Sciences. The pseudostem of this plant is about 3.5m tall. (B) Inflorescence with male and female flowers showing bracts and flowers alternately arranged along the main axis. (C) Staminate flowers, and visiting black shield wasp (*Vespa bicolor,* Vespidae, Hymenoptera) (D) Female flowers. (E) Fruits. Bars in B-E represent 2 cm. (F) Native distribution of *Ensete glaucum, E. superbum, Musa* and *Ensete* species by countries or provinces (for China, India-Assam, and Australia). Musaceae are not currently native in the Americas, although *Ensete* is present in the fossil record (Manchester et al., 1993). *E. glaucum* always occurs in the same provinces as *Musa* and sometimes with other Asian *Ensete* species. Map adapted from POWO [52].

Originating in the tropics and subtropics at lower elevations, most species in Musaceae lack cold acclimation. Cold stress is one of the key limitations in extending banana planting and production to higher altitudes and beyond the tropics [12]. In contrast to other Musaceae species which would not survive, *Ensete glaucum* can be found above 1000m in the mountains of Yunnan in China, where the temperature drops lower than 0°C, with limited rainfall in winter. As one of the most cold-resistant and perhaps the most drought-tolerant species in Musaceae, *E. glaucum* is a potential gene and germplasm resource for abiotic stress tolerance in banana breeding, likely to be required for the adaptation to a more variable and extreme climate in the future.

Whole genome sequences (genome assemblies) are published for some species in Musaceae, with pseudochromosome-level data only available in *Musa* [13–15]. Draft genome assemblies in *Ensete* species are limited to only *Ensete ventricosum,* and these are with tens of thousands of contigs with an N50 lengths mostly between 10,000 and 21,000 bp [16, 17]. As such, effective introduction and utilization of genetic resources present in wild species of *Ensete* are a pressing need for banana improvement.

We applied Illumina, Oxford Nanopore Technologies (ONT), and chromosome conformation capture (Hi-C) sequencing to generate a high-quality chromosome-level assembly of the *Ensete glaucum* genome. We aimed to use the chromosome sequence to show the genome structure and gene composition, as well as revealing the repetitive DNA organization. The structural variations of *Ensete glaucum* (*x* = 9) were studied in a comparative context with *Musa* (x = 11) species, showing the evolutionary history of the family. The study aims to be useful in expanding the genepool available not only to banana and enset breeders, but also for plant conservation of biodiversity in ecologically sensitive or threatened areas, and for fundamental research on chromosome and genome evolution.

## Analyses, Results and Discussion

### *De novo* chromosome-scale genome assembly

A *de novo* chromosome level assembly of *Ensete glaucum* was made by combining high-coverage ONT long-read sequencing, Illumina 150bp paired-end sequences, and Hi-C chromosome conformation capture sequence data (Table 1). From the first assembly (with 124 contigs with an N50 of 10.256 Mb, Table 2 and Supplementary Table S1), we assembled nine pseudomolecules, eg01 to eg09 (Fig. 2 and Supplementary Table S2), corresponding to the chromosome number (2n=18) and observed chromosome morphology.

**Figure 2.**
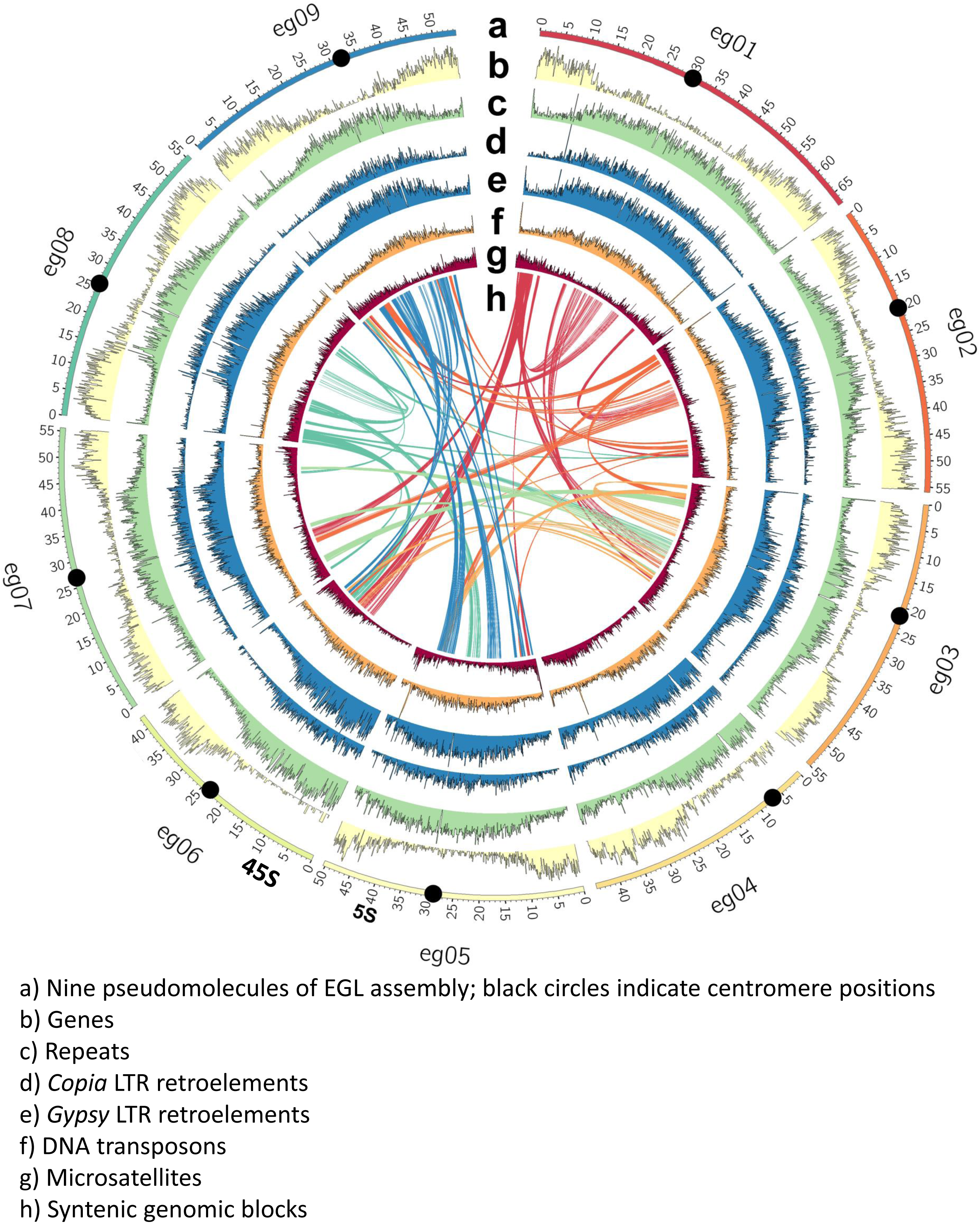
*Ensete glaucum* chromosome assembly and genome features. Circos plot of (a) The nine pseudomolecules of the EGL assembly corresponding to 9 chromosomes. Black dots indicate centromere positions and 5S and 45S the positions of rDNA loci; scale in Mbp; (b) Gene density; (c) Repeat density; (d) *Copia* LTR retroelement density; (e) *Gypsy* LTR retroelement density; (f) DNA transposon density; (g) Short tandem repeat (microsatellite) density; (h) Syntenic genomic blocks, linked by curved lines (arbitrary color) in middle of the plot.

**Table 1:** Statistics of whole-genome sequencing and transcriptome analysis of *Ensete glaucum* using Illumina, ONT, and Hi-C.

| Type | Method | Number of reads | Clean data (Gb) | Length (bp) | Coverage (×) |
| --- | --- | --- | --- | --- | --- |
| Genome | Illumina | 245,852,534 | 37 | 2 × 150 | 74 |
|  | ONT | 4,357,035 | 109 | 38,885 (N50) | 220 |
|  | Hi-C | 319,793,734 | 48 | 2 × 150 |  |
| Transcriptome | Illumina (Leaf) | 62,410,840 | 94 | 2 × 150 |  |
|  | Illumina (Root) | 13,712,542 | 21 | 2 × 150 |  |

**Table 2:**
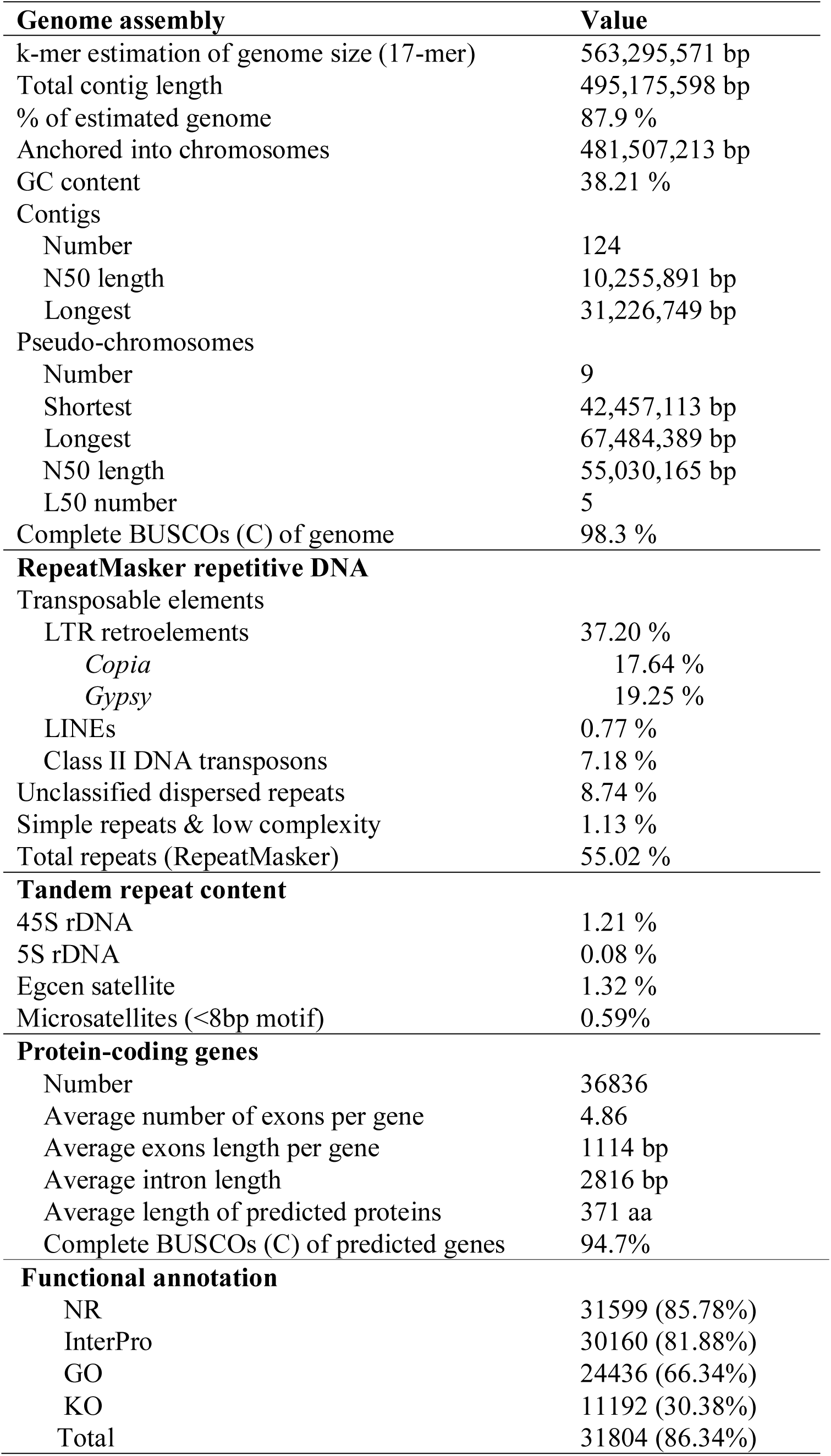
Statistics of *Ensete glaucum* genome assembly and annotation. RepeatMasker did not identify satellite sequences. 5S, 45S and Egcen were identified manually in assemblies and the abundance measured in raw read data; microsatellite abundance was calculated from the assemblies. (See Supplementary Tables S1, S2, S3, S10, S12, S14)

A Hi-C/ONT-only assembly was constructed first, by using an OLC (overlap layout-consensus)/string graph method with corrected reads. Contigs were refined using Illumina short reads, and after discarding redundant contigs, the final genome assembly was 481Mb long, with 9 chromosomes between 42,457,113 and 67,484,389bp long. Our BUSCO analysis [18] was used to assess the assembly in ‘genome’ mode showing 98.3% complete single and duplicated Embryophyta core gene sets from the embryophyta_odb10 database (consisting of 1614 genes) (Table 2, Supplementary Table S3a); few genes were fragmented or missing. *E. glaucum* chromosome designations were chosen to follow major regions of synteny with *Musa acuminata* chromosomes [19, 20].

### Genome size and heterozygosity estimation

The final genome assembly after Hi-C scaffolding (481,507,213 bp) anchored 97.2% sequences of the contig-level assembly (495,175,598 bp; Table 2). Some arrays of tandem repeats, including the rDNA (see below), were collapsed and chromosome termini were not fully assembled. Around 55% of the assembled genome was estimated to be repeat sequences (RepeatMasker; Table 2). The genome size was estimated as 563,295,571bp (highest 17-mer peak frequency), with slightly higher estimates made by findGSE software (588,939,614 bp). The genome size of *E. glaucum* is similar to that of the *x*=11 *Musa* species (see [20]) using sequencing methods, and to estimates of both genera by flow cytometry [21].

The heterozygosity rate of *E. glaucum* was 0.164% (Supplementary Fig. S1 estimated with k=21 using GenomeScope). Heterozygosity in plants is influenced by mating-systems and pollination [22], life span, habitat fragmentation and cultivation [23]. There is relatively little known about the breeding system and pollination of *Ensete* species (see [24]) although we observed insects (including the hornet *Vespa bicolor,* a widespread pollinator in southern China) visiting flowers (Fig. 1C). Our low level of heterozygosity is within the range found in individual plants in populations of *M. acuminata* ssp. *banksii* (0.02% to 0.34% in 24 individuals, [25]; and 0.13% to 0.23%, [26]), and in other wild monocotyledonous species including two (most likely self-pollinating) diploid oat species (0.07% heterozygosity in *Avena atlantica* and 0.12% *A. eriantha* [27]); it is however, low compared to other species (e.g. walnut, *Juglans nigra* 1.0%, [28]; *Nyssa sinensis* 0.87%, [29]) and in particular many *Musa* species, some with known hybrid genome composition [26]. The low value seen in species including *E. glaucum* here, is consistent with frequent self-pollination and inbreeding, or a population bottleneck of this monocarpic tropical plant [30].

### Gene distribution and whole-genome duplication analysis

Genes were unevenly distributed along chromosomes (Fig. 2 circle b), and generally depleted in broad centromeric regions; few genes were found on the short arm of the more acrocentric chromosome eg04 and the Nucleolus Organizing Regions (NOR) bearing chromosome arm of eg06 (see rDNA below). The centromeric, gene poor regions, are rich in repeats (Fig. 2 circle c), transposable elements (*Copia* and *Gypsy* LTR retroelements, Fig. 2 circles d, e, and, less markedly, DNA transposons, Fig. 2 circle f), as observed in many species (including *Musa*,[14, 15]).

The *K*s (the synonymous rates of substitution) between genes in paired collinearity gene groups were calculated between *E. glaucum* and *M. acuminata* to see whether they share the same three WGD events [14]. The two genomes have a nearly identical *K*s density distribution (Fig. 3A), both having two peaks at around 0.55 and 0.9. This result indicates that Musaceae share the same WGD events. The more recent peak at 0.55 most likely represents the α and β duplications, while the peaks at 0.9 may represent the more ancient γ duplication event [14]. Fig. 2 (center) links the genomic locations of paralogous gene clusters: most chromosome regions show shared relationships with three other chromosome regions, reflecting the α and β whole genome duplications, as shown by D’Hont et al. [14] (their Supplementary Figure 12).

**Figure 3.**
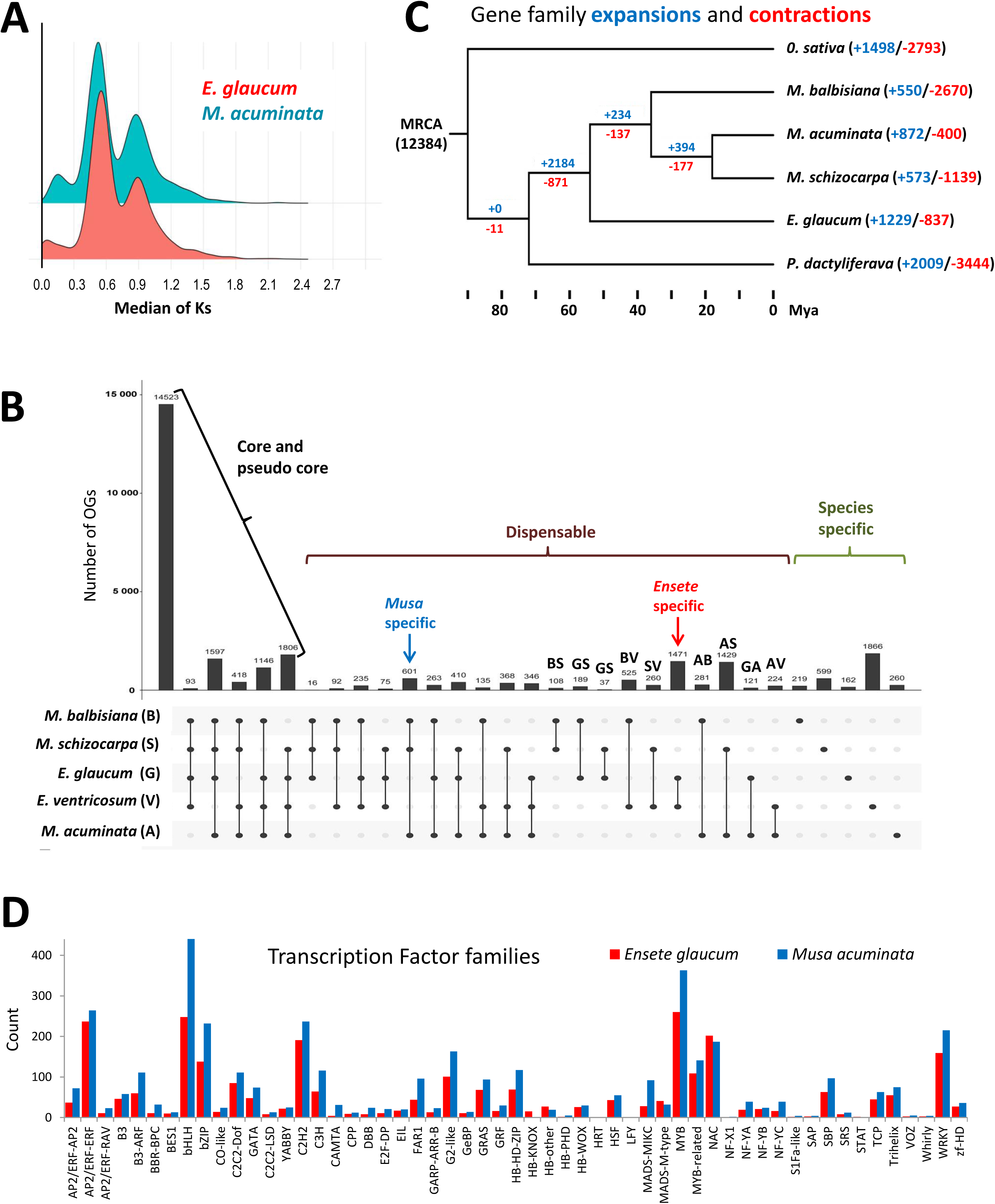
Gene family evolution and conservation. (A) Gene family expansion and contraction with a phylogenetic tree showing timeline of divergence of monocot species. MRCA: Most Recent Common Ancestor. Numbers denote the gene family expansion (orange) and contraction (green). (B) The synonymous substitutions (*K*s) frequency density distributions of orthologs within EGL or MAC, whose peaks indicate whole-genome duplications (WGDs). (C) Intersection diagram showing the distribution of shared orthogroups (OGs) (at least two sequences per OG) among *Musa* and *Ensete* genomes. codes: E, *E. glaucum*; V, *E. ventricosum;* A, *M. acuminata*; B, *M. balbisiana*; S, *M. schizocarpa.* (D) Histogram of the comparative abundance (umber of of transcriptions factors between *M. acuminata* and *E. glaucum.*)

### Gene identification

#### Genes and gene ontology

In total, 36,836 genes were predicted (BUSCO score: C:94.7%, Supplementary Table S3B) with 31,804 (86.34%) functionally annotated with protein domain signatures and 24,436 (66.34%) associated with GO terms (Table 2; Supplementary Table S4). *E. glaucum* has a similar gene space (Supplementary Table S5) to the sequenced *Musa* species *M. acuminata* (35,264), *M. balbisiana* (35,148), *M. itinerans* (32,456) and *M. schizocarpa* (32,809). In the Musaceae (i.e., *M. acuminata, M. balbisiana, M. schizocarpa, E. ventricosum* and *E. glaucum*), we identified a total of 29,639 orthogroups including 173,025 (88.1%) assigned genes and 23,355 (11.9%) unassigned genes (Fig. 3B and Supplementary Table S5). Between all species, the analysis showed 48% (n=14,523) of assigned genes were shared (core or pseudo-core genes; rising to 66%, n=19,583 if genes missing in only one species are discounted as possible annotation artefacts). The analyses highlighted 5% (1471) of orthogroups that are *Ensete* genus specific and not found in *Musa*. A total of 162 orthogroups were found only in *E. glaucum* (Fig. 3B; lower than the value for *E. ventricosum* but the latter is a draft genome status without RNA support and with fragmented contigs with likelihood of a large number of redundant predicted genes). The predicted genes of *Ensete glaucum* were compared to their orthologous genes in *M. acuminata* and *K*a/*K*s values between orthologous pairs were calculated. Genes with *K*a/*K*s > 1 were under positive selection (Supplementary Table S6).

#### Gene family expansion and contraction

Using *Musa* species and two other monocotyledonous species (in the same clade of the Commelinids), *Phoenix dactylifera* (Arecaceae) and *Oryza sativa* (Poaceae), we explored gene family expansion and contractions in *E. glaucum* (Fig. 3C and Supplementary Table S7). Among 12,384 gene families shared by the MRCA (Most Recent Common Ancestor) of these monocotyledons, there were large numbers of gene families expanding (1498 to 2184) or contracting (817 to 3444) between the genomes of Musaceae, *Phoenix* and *Oryza* (Fig. 3C), presumably reflecting substantial differences in plant form between them. Similar, although slightly lower, figures were reported between, for example, dicotyledons as diverse as *Arabidopsis* (Brassicaceae), *Solanum* (Solanaceae) and *Cuscuta* (Convolvulaceae) [31]. Notably, though, our results show the largest expansion of gene families in the Musaceae (2184), likely reflecting the whole genome duplication events not shared with the Poaceae or Arecaceae (see also [32] in pineapple), and we find additional expansion in *E. glaucum*. Large gene family losses were noted in *Oryza sativa*, *Phoenix dactylifera* and *Musa balbisiana* (Supplementary Table S7).

Overall, *E. glaucum* showed enrichment of several GO biological processes (Supplementary Fig. S2, Supplementary Table S8) compared to *Musa*. Among them, “monosaccharide transmembrane transporter” (equal top hit), “carbohydrate transmembrane transport” and “carbohydrate transport”; and among molecular functions, “monosaccharide transmembrane transporter activity”, “sugar transmembrane transporter activity”, and “carbohydrate transmembrane transporter activity” were all included in the top 20 enrichments. The genus *Ensete* is notable for its accumulation of starch in the pseudostem and leaf bases with *E. ventricosum* cultivated as a staple starchy food in East Africa [2], and perhaps this is reflected in the enrichment of certain carbohydrate transport GO terms.

#### Transcription Factors (TFs)

In total, 2637 putative TF genes were identified in the *E. glaucum* assembly, representing 7% of all genes (Supplementary Table S9), which were classified by their signature DNA Binding Domain (DBD) into 58 TF families (Fig. 3D). Similar to *M. acuminata*, the MYB (myeloblastosis) superfamily of transcription factors (including 260 MYB TFs plus 109 MYB-related) was the largest family, with between 140 and 210 copies of each of the bHLH, AP2/ERF, NAC, C2H2, WRKY and bZIP families. The identification and classification of the TFs here provides a framework to explore regulatory networks in plants [33] with their target genes, and to identify specific factors involved in important responses. Cenci et al. [34] analyzed transcription factors involved in the regulation of tissue development and responses to biotic and abiotic stresses and, particularly, the NAC plant-specific gene family, while Xiao et al [35] discuss the importance of a HLH factor involved in starch degradation during fruit ripening. *Ensete* and *Musa* differ in these characteristics so it will be interesting to analyze differences in transcription factors responsible.

### Repetitive DNA analysis

#### Repeat identification

A range of different programs were applied for repeat analysis, and, as has been considered previously [36], there were differences in the repeats identified between approaches, and small changes in parameters and reference sequences give substantial changes. Repeated elements in the genome assembly were identified by RepeatMasker (Table 2 and Supplementary Table S10) and amounted to 55% of the genome, the same range as other plant species with similar DNA amount and, particularly, the genus *Musa* [14, 15]. For assembly-free identification of repeats, we used RepeatExplorer [37] to generate graph-based clusters of similar sequence fragments: Illumina sequence reads are available from six Musaceae species, allowing assembly-free comparisons (Supplementary Table S11 and Supplementary Fig. S3); while there was a little more variation in proportion of reads in the most abundant clusters, all had between 33% and 46% in the top clusters (>0.01% genomic abundance, as defined in [35]). Transposable elements including LTR and non-LTR retroelements, and class II DNA transposons, were found (Fig 4 and Supplementary Table S11). Microsatellites and other repeats were further characterized by mining and dotplot analysis as well as fluorescent *in situ* hybridization to chromosomes (Supplementary Tables S12-S14 and Figs 5-7; see below). The organization of repetitive regions in the assembly was sometimes verified by mapping individual ONT long reads to assembled repeat regions (eg. Fig. 5A), and organization was generally confirmed, except for some long tandem arrays which seem to be collapsed in the assembly due to high homology between repeat units. A few ONT reads were found which included reversals of tandem arrays (head-to-head or tail-to-tail junctions), potentially artefacts from both strands of the DNA molecule passing sequentially through one pore, and these junctions need further investigation.

**Figure 4.**
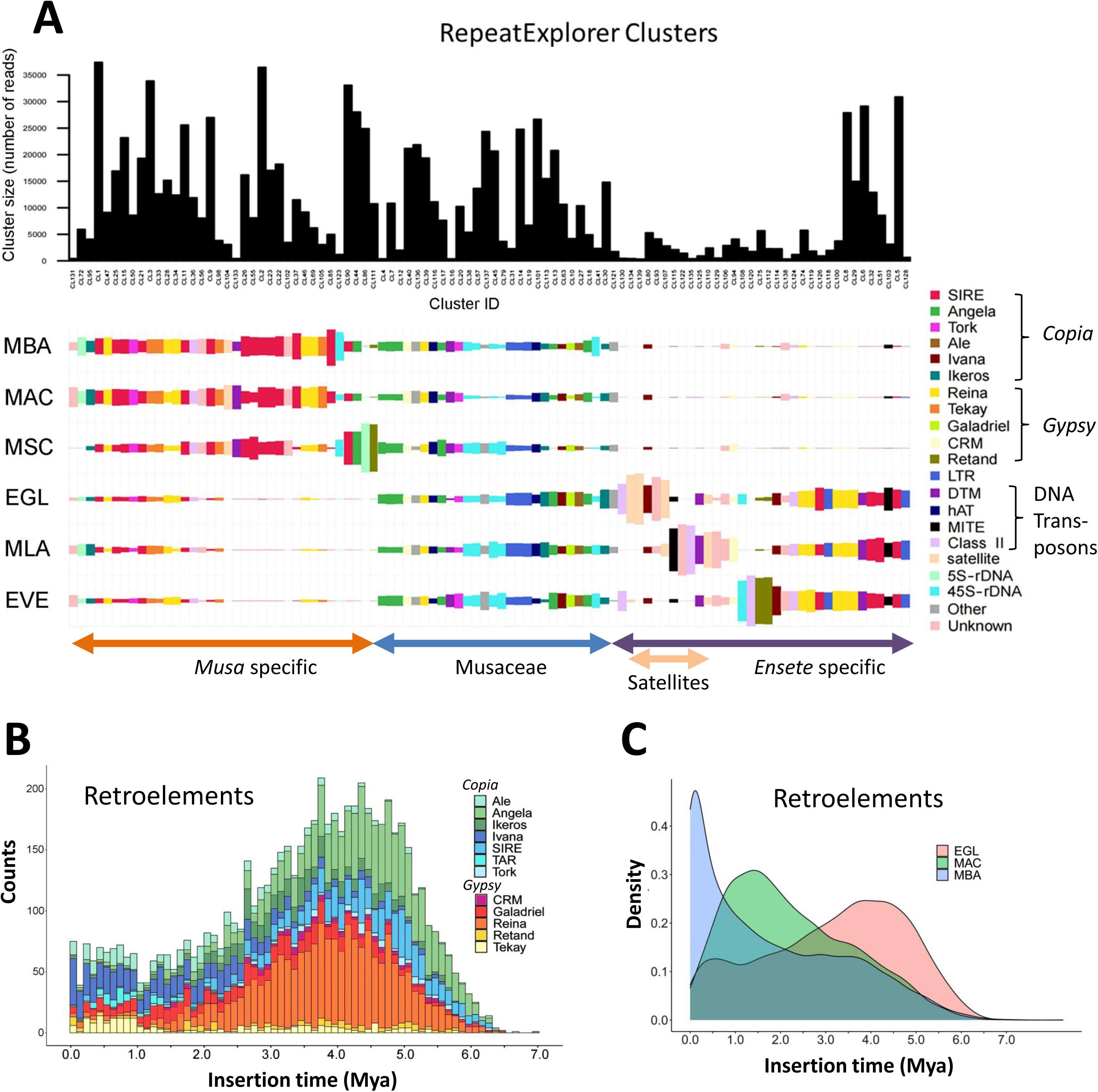
Comparative analysis of repetitive DNA in Musaceae using RepeatExplorer. (A) Bar plot showing the sizes (numbers of reads) of the most abundant individual graph-based read clusters with (lower part) distribution among six Musaceae species (coloured rectangle size is proportional to the number of reads in a cluster for each species. Clusters and species were sorted by using hierarchical clustering. Individual rectangles are colored based on the annotation of the clusters. Figure was adapted from the output of the visualization script (plot_comparative_clustering_summary.R). Species codes: EGL, *Ensete glaucum*; EVE, *E. ventricosum*; MAC, *Musa acuminata*; MBA, *M. balbisiana*; MLA, *Musella lasiocarpa*; MSC, *M. schizocarpa*. (B) The nature and ages of LTR retroelements in *E. glaucum*. (B) The ages of retroelement insertions in *E. glaucum, M. acuminata* and *M. balbisiana*. Mya = Million years ago.

**Figure 5.**
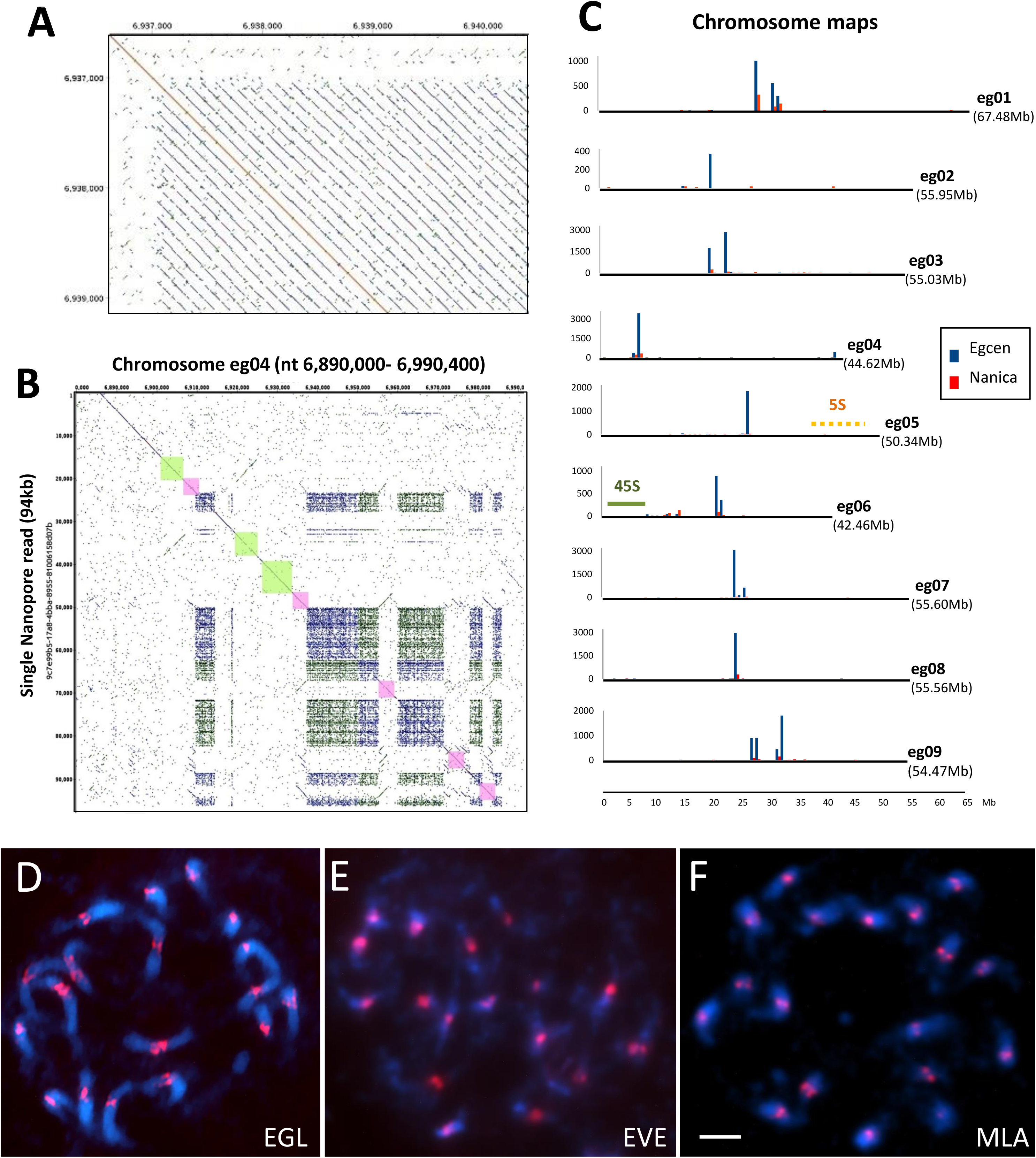
*Ensete glaucum* centromeric repeat structure. (A) Dot plot (self-comparison of sequences) showing start of a 134bp Egcen tandem array. (B) Dot plot showing part of a chromosomes assembly (eg04) plotted against part of a single ONT read with blocks of the Egcen tandem repeat (appearing as dense rectangles at this scale) interspersed with Nanica elements (red; five homologous copies in both orientations) and LTR retroelements (green; two non-homologous sub-families), (C) Bar chart showing frequency distribution of the EGcen centromeric tandem repeat, Nanica transposable elements (x10 on axis), and locations of 45S and 5S rDNA along the assemblies for each chromosome. Long Egcen arrays occur at one of more sites at the centromeric regions of all chromosomes. (D, E, F) *In situ* hybridization of Egcen probes detected by red fluorescence to cyan-fluorescing DAPI-stained chromosomes of (D) EGL, *Ensete glaucum*; (E) EVE, *E. ventricosum*; and (F) MLA, *Musella lasiocarpa.* The red Egcen locations collocate with the primary centromeric constriction on all 9 pairs of chromosomes. Bar=5µm.

**Figure 6.**
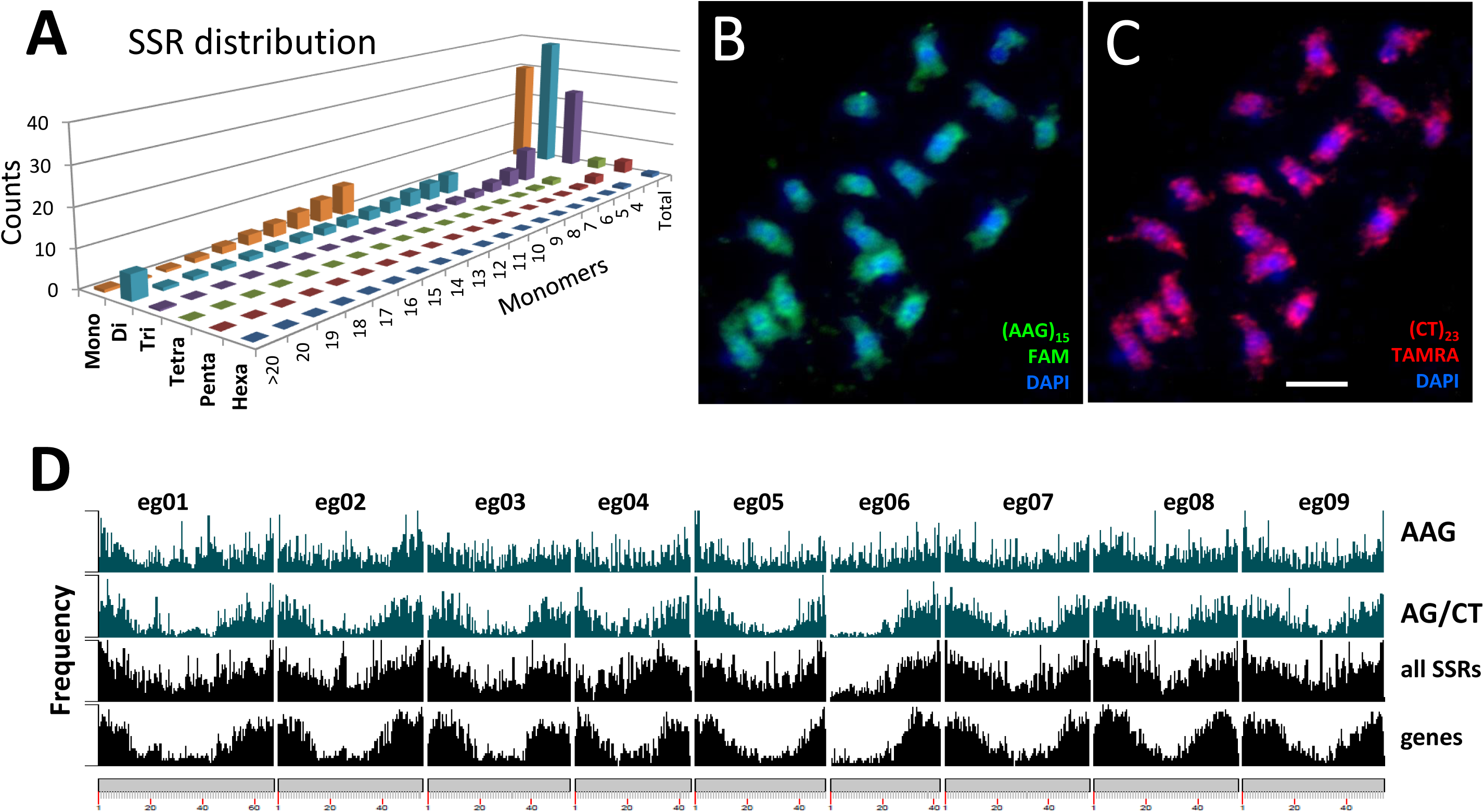
Microsatellite (SSR) distribution in *Ensete glaucum*. (A) Abundance (count) and total number of monomers microsatellites (SSR) with motifs between 1 and 6 bp long. (B, C) *In situ* hybridization of synthetic microsatellite probes to DAPI-stained (blue) chromosomes, showing (B) AAG is relatively uniformly distributed along chromosomes compared to (C) where the greater abundance of AG/CT in distal chromosome regions is seen. (D) Abundance of AAG, AG, all microsatellites, and genes along the chromosome assemblies. In agreement with the *in situ* hybridization result, AAG is more uniformly distributed, while AG (along with genes and all the microsatellites pooled) show greater abundance in distal chromosome regions except for the arm of chromosome eg06 carrying the 45S rDNA (NOR). Bar=5µm.

**Figure 7.**
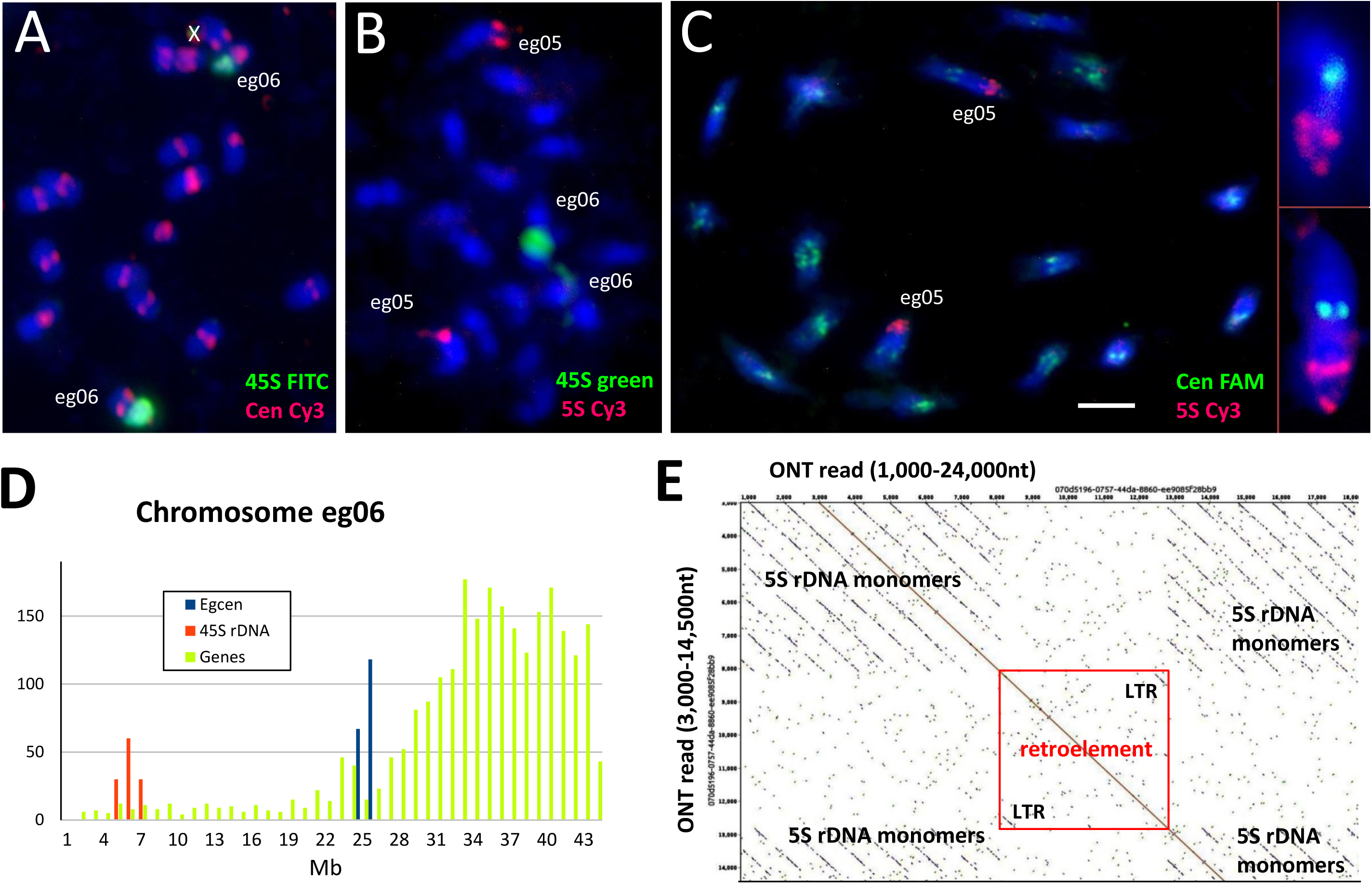
rDNA in *Ensete glaucum*. (A-C) *In situ* hybridization to chromosomes (stained blue with DAPI) showing locations of (A) the 45S rDNA (green) on one pair of chromosomes while Egcen (red) locates all 9 pairs of Chromosomes showing location of 45S (green on eg06; the two sites are on a pair of chromosomes which are adjacent to each other) and 5S the 5S rDNA (red) in a more dispersed pattern on one pair of chromosomes with Egcen (green) at all 9 pairs of centromeres; inset shows 5S rDNA chromosome pair at higher magnification. The 5S rDNA signal is dispersed over a longer region of a chromosome, while the 45S locus occupies much of the chromosome arm. Bar 5µm. X=stain precipitate. (D) Histogram showing density of genes (light green), Egcen (blue) and 45S rDNA copies on chromosome eg06. The arm carrying the 45S rDNA is depleted in genes. (E) Part of a single ONT read covering 24kb spanning part of the 5S rDNA array. The unusually long 1056bp tandemly repeated 5S rDNA monomers (14 copies) are interrupted by an LTR retroelement. LTRs, with no homology to the 5S rDNA, are seen (bottom left and top right) in the red box.

Fig. 4A compares the abundance and species-distributions of major repeat classes in the Musaceae using the comparative genome analysis function of RepeatExplorer. All species shared many transposons and rDNA sequences (Fig 4A, central region). However, genus-specific retroelement variants were identified in *Musa* (Fig. 4A, left) and *Ensete*-with- *Musella* (Fig. 4A, right), showing the separation of the two phylogenetic branches, supported by extensive divergence of the repetitive sequence sub-families, and evolution in copy number. Notably, satellite sequences (Fig. 4A center-right) were much more abundant and some sequences (see centromere sequence below) were present exclusively in *Ensete*.

#### Transposable Elements

The most abundant class of repetitive elements were transposable elements, in particular LTR retroelements. The distributions of *Copia* and *Gypsy* LTR retroelements along assembled pseudo-chromosomes (Fig. 2 circles d and e) show greater abundance in proximal chromosome regions. Approximately equal numbers of *Copia* and *Gypsy* elements (18 and 19% of the genome respectively, Table 2) were found. This result contrasts with *M. acuminata,* where *Copia* elements were considerably more frequent (29%) compared to *Gypsy* elements (11%;[14]; Supplementary Table S11). The relative change in proportions of the two element families, while the overall abundance remains the same, has implications for genome evolution and the expansion or contraction of retrotransposon families which can be explored in detail using the high-quality genome sequences where the elements are neither truncated nor collapsed.

Analysis of RT domains identified subfamilies of LTR retroelements, with the families showing different abundances in *E. glaucum* and *M. acuminata* (Supplementary Fig. S4). Insertion times of LTR retroelement subfamilies (Figs 4B, C and Supplementary Fig. S5) were calculated based on LTR divergence for *E. glaucum* and recalculated for *Musa* to allow for identical software settings (see Material and Methods). In *E. glaucum,* both *Copia* and *Gypsy* families show relatively constant activity over the last 2.5 My, with the major a peak of insertion activity 3.5 to 5.5 Mya, (Fig. 4B, C) corresponding to the half-life of LTR-elements [14]. The dynamic amplification of these elements is emphasized by individual sub-families having bursts of amplification (Fig. 4B for *E. glaucum* and Supplementary Fig. S4 for *Musa*), with rounds of expansion of different elements. As shown by Wang et al. [15], *M. balbisiana* has the most extensive LTR activity in the last 500,000 years, and *M. acuminata* activity peaks around 1.5 Mya (Fig. 4C), in both cases with greater activity of *Copia* elements (Supplementary Fig. S5), contrasting with *E. glaucum* with equal activity of both *Gypsy* and *Copia* elements leading to a higher proportion of *Gypsy* elements within the genome of *E. glaucum* compared to *Musa* (see above, and Supplementary Tables S10 and S11). This is also evidenced by the larger number of *Musa* specific clusters identified as *Copia* Angela or Sire elements while *Ensete* with *Musella* specific LTRs include more *Gypsy* Reina and Retand elements (Fig. 4A). Wu et al. [38] discuss the rounds of amplification in *M. itinerans* with an amplification burst after separation from *M. acuminata* about 5.8 Mya suggesting high turnover of the elements. The results suggest a burst of retroelement amplification (the older ones), sometime after the split of *Musa* and *Ensete,* and again more recently, perhaps after *E. glaucum* split from other *Ensete* species.

#### Tandem (satellite) repeats and centromeric sequences

The repeat analysis revealed the presence of an abundant tandemly repeated sequence with a monomer length of c. 134bp (Fig. 5A). The sequence, named Egcen, represents about 1.3% of the *E. glaucum* genome (45,000 copies) (Table 2, Supplementary Table S12; GenBank: OL310717), and forms arrays that are at places interspersed by the LINE element Nanica (described in *M. acuminata;* [14]) and other sequences (Fig. 5B, see below). One or two major arrays of Egcen repeats were found in the assemblies of all nine chromosomes (Fig. 5C). *In situ* hybridization of Egcen showed it was located around the primary, centromeric, constrictions as seen by DAPI staining (Fig 5D, see also Figs. 7A and C). The hybridization pattern of the FISH signal on all chromosomes showed variable strength and several sites grouped closely together, corresponding to the pattern seen in the assembly. The location of the Egcen arrays was therefore used to infer the centromere mid-point position in the *E. glaucum* chromosome assemblies (Fig 2 outer circle, Fig. 8A and Supplementary Table S13).

**Figure 8.**
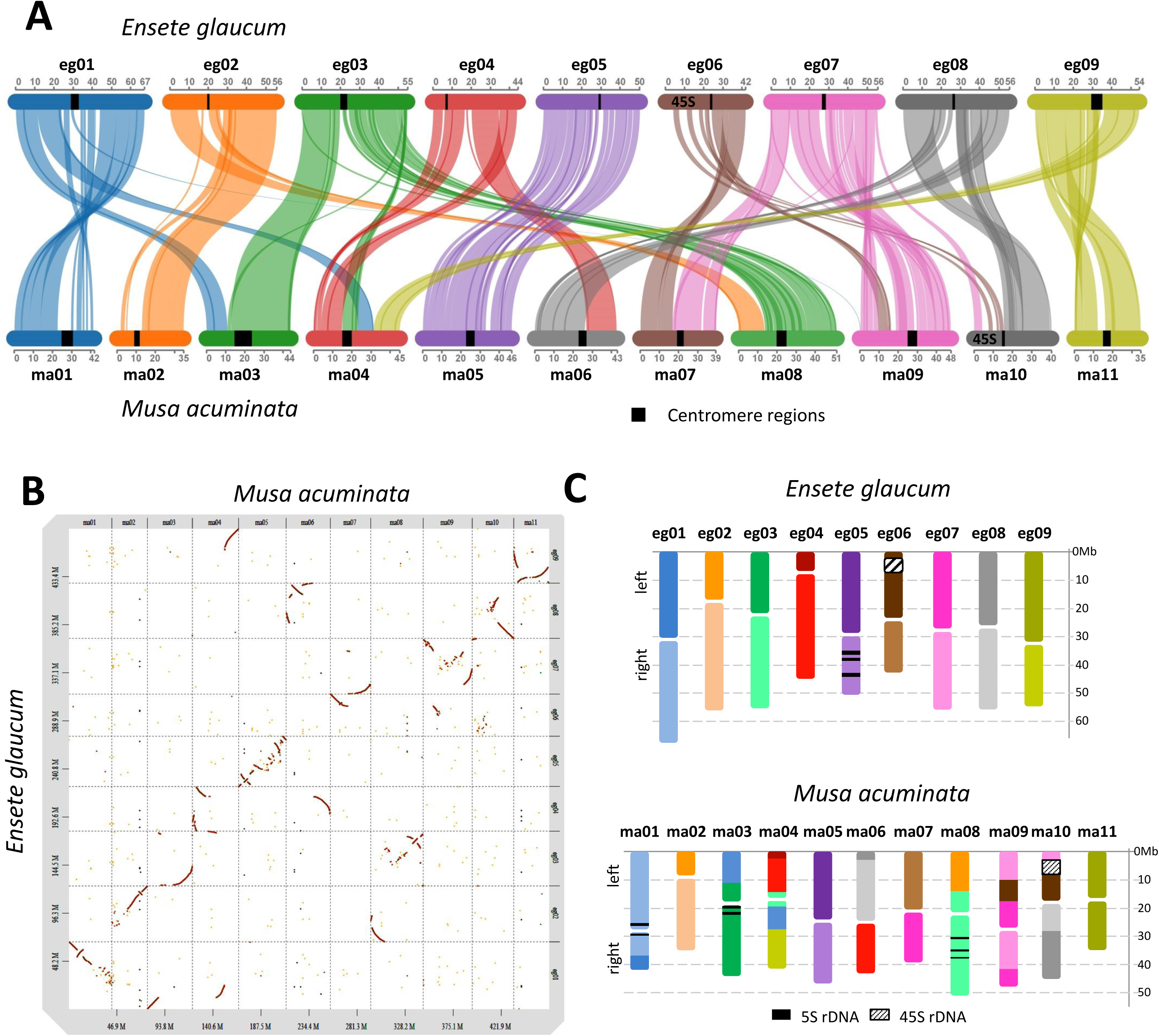
Synteny of Ensete glaucum and Musa acuminata. (A) Synteny plot Synvisio connecting syntenic genes in the 9 chromosomes of *E. glaucum* (egxx) and 11 chromosomes of *M. acuminata* (maxx). syntenic blocks of high homology are indicated by uniformly coloured areas in the graphs. Only eg05 and ma05 maintain synteny over the full chromosome length, although there are some rearrangements. Three ma chromosomes are represented by part of one eg chromosome, while other ma chromosomes are fusions of more than one eg chromosome. B) Dot plot comparing DNA sequences of *E. glaucum* and *M. acuminata*. C) representation of the syntenic blocks in the karyotypes of *E. glaucum* and *M. acuminata*. Chromosome rearrangements are shown, complementing the Synteny plot, while inversions and relative expansions and contractions of genome regions are clear.

Egcen was also detected at the centromeres of all *E. ventricosum* and *Musella lasiocarpa* chromosomes (Fig. 5E, F), showing similar distribution patterns as in *E. glaucum* (Fig. 5D), but it was not seen on *Musa* chromosomes by *in situ* hybridization (example of *M. balbisiana,* Supplementary Fig. S6) nor found in analysis of assemblies of *M. acuminata, M. balbisiana* or *M. schizocarpa* (Supplementary Fig. S7A). The comparative RepeatExplorer clustering shows multiple satellite sequences found only in the *Ensete* and *Musella* genomes that are not present in the three *Musa* species tested (Fig. 4A) and supports Egcen being part of the tandem repeat birth and amplification that has occurred in *Ensete* and *Musella* after the split from *Musa,* and contrasts with the younger insertion times found for *Musa* retroelements (Supplementary Table S12 and Fig. 4C).

Tandem repeats or satellite DNA sequences are found around the centromeres of many plant (and animal) species [39, 40] and may be ‘centromeric’ or ‘pericentromeric’. No equivalent tandem repeats were found in *Musa* [14,20,41] and the *E. glaucum* Egcen is not present either in *Musa* (Supplementary Figs S6 and, S7A). However, the centromeric regions of all *M. acuminata* chromosomes have been shown to include multiple copies of a LINE non-LTR retroelement, Nanica, both by *in situ* hybridization and bioinformatic analysis [14, 20]. Nanica-related sequences were also identified in the *E. glaucum* assembly, but with less abundance than in *Musa* (Supplementary Fig. S7B); about 350 copies were mapped to chromosomes, mostly (but not exclusively) present interspersed within, or adjacent, to Egcen arrays (Fig. 5B, C and Supplementary Fig. S7B).

Assembly across centromeric regions including abundant repeats is difficult and normally the tandem repeat elements are collapsed. The ONT long-molecule sequences allowed detailed examination of parts of the centromere region of chromosomes. A dotplot of an ONT read (coded 9c7e99b5, 96,300bp long) aligned to the assembly of eg04 shows the complex organization of the Egcen array (Fig. 5B): this 100kb region includes a total of six Egcen tandem blocks with between 3 and 126 repeats (a total of 385), five copies of Nanica (some rearranged, pink boxes), and three diverse retroelements flanked by LTRs (green boxes). Further copies of the Egcen tandem repeat occur in larger blocks over the following 450,000bp of the assembly, and no genes were identified in the region.

A characteristic 17bp long sequence, the canonical CENP-B box, is found within a monomer of a tandem repeat at centromeres of many species including human [42] and has been postulated to be necessary for binding of the centromeric CENP-B proteins regulating formation of centromere-specific chromatin. Within the Egcen sequence, there was a CENP-B related motif:

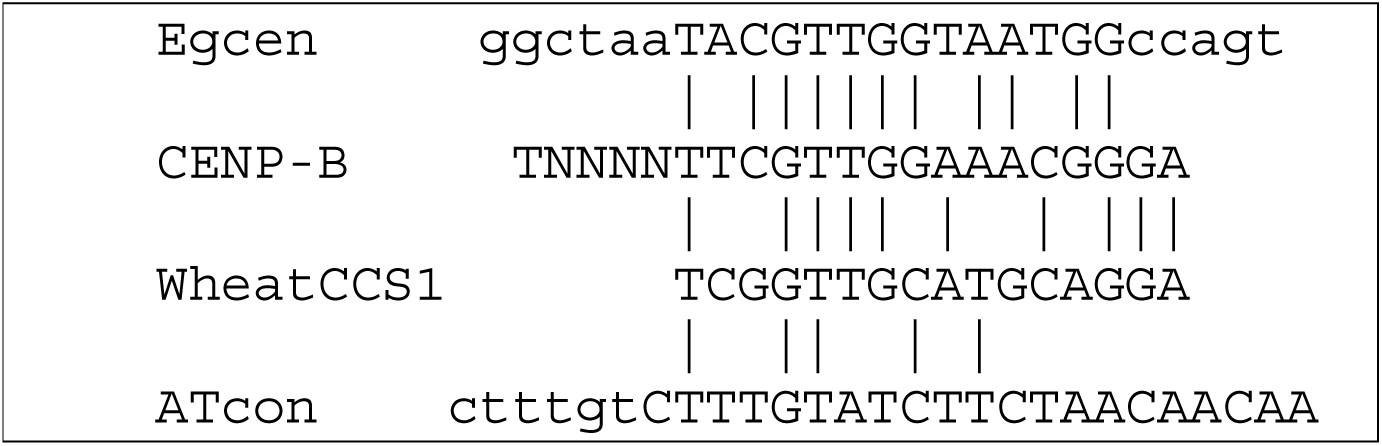

Although, similar CENP-B motifs have been found from wheat and *Brachypodium* (CCS1, [43]) to *Arabidopsis* (ATcon, [44]; see also review [45]), the relevance of the CENP-B related box to centromere function remains uncertain, particularly when no similar tandem repeat is present in other species or the related genera such as *Musa* (see above). As is the case in many other species, the relative roles of retroelements, tandem repeats, and interspersed centromeric sequences, leading to recruitment of the centromeric proteins, are uncertain: the exact sequence or sequences that mark the functional centromeres, remain enigmatic. The identification of a centromeric tandem repeat and the assembly across all centromere regions in *E. glaucum* together with data from *Musa* will allow protein binding studies (with Chromatin Immunoprecipitation ChIP analysis) to resolve the functional centromere.

#### Microsatellites (Simple sequence repeats, SSRs)

Microsatellites were searched using Phobos and an SSR mining pipeline ([46]; perfect SSRs from mono to hexa-nucleotide repeats with 11 to 3 repeat numbers respectively). SSR abundances, array lengths, and nucleotide base composition (70% were AT-rich) are shown in Fig. 6A and Supplementary Table S14. The overall nature and abundances of microsatellites were generally similar to *M. acuminata*, *M. balbisiana, M. itinerans* and *E. ventricosum* (Table S14). An average of one SSR was found per 4000bp, with the density lowest around the centromere and higher at the telomeres (Fig. 6B); they are excluded from the 45S NOR chromosome arm of eg06, and their overall distribution is similar to the distribution of genes but contrasts the more proximal distribution of LTR retroelements and DNA transposons (Fig. 2). Individual microsatellite motifs, however, showed characteristic and different distributions. The abundant microsatellites, CT and AAG, were synthesized as labelled oligonucleotides probes and used as probes for FISH on chromosomes. Both the bioinformatic analysis of the assembly (Fig. 6B) and FISH to chromosomes (Fig. 6C, D) showed (AAG/CTT) has a relatively uniform distribution along chromosomes, while (AG/CT) shows depletion in centromeric regions and greater abundance in distal parts of chromosomes that are gene-rich (Fig.2). The constraints on microsatellite spread in the genome are motif-specific, and, if SSR markers were to be used for genetic mapping, those associated with genes (such as AG/CT) would potentially be more useful.

#### 5S and 45S rDNA and rRNA genes

Tandem repeats of the rDNA were predominantly located within extended, complex, loci on chromosomes eg05 (5S rDNA) and eg06 (45S rDNA) (Figs 2, 5C, 7, and 8). The 45S rDNA monomer containing the 18S rRNA gene - ITS1 - 5.8S rRNA gene - ITS2 - 26S rRNA gene - NTS (GenBank: OL310719) was 9984 bp long, typical but slightly longer than other plant species. The NTS region includes in most cases 16 copies of a degenerate 180bp tandem repeat. Based on occurrence in the unassembled Illumina reads, there were 587 copies of the 45S rDNA monomer (1.21% of the genome, Tables 2 and Supplementary Table S12). Although in the whole genome assembly the rDNA array was collapsed, the strength of the *in situ* hybridization signal using the rDNA sequence from wheat (Figs 7A,B) is consistent with representing 1% of the genome. The long chromosome arm carrying the 45S NOR locus was depleted in genes by 10-fold (average of 12.6 genes/Mb compared to 127.6/Mb on the short arm; (Fig. 7D). The single site of 45S rDNA at the NOR (Nucleolar Organizing Region) per chromosome set is similar to *Musa acuminata* and other *Musa* species [47], although not *E. gilletii* (2n=18), where there are 4 pairs of sites [21].

The 5S rDNA (GenBank: OL310718) comprised the 5S rRNA gene (119bp long, typical for all plants) and intergenic spacer (937bp), representing 0.078% of the genome or approximately 366 copies, with a complete motif length of 1056bp (Supplementary Table S12). The 5S rDNA locus lies in the middle of the short arm of chromosome eg05, in three parts around 34.5M, 37.5M and 45.5M with multiple interruptions. An example of insertion of a 4.7kb LTR retrotransposon-related sequences in the 5S rDNA array of tandem repeats is shown in Fig 7E. In other regions of the *E. glaucum* ONT reads or assembly, the retroelement-related sequences named Brep, reported in *Musa* [48], was also found in the 5S rDNA arrays. Garcia et al. [49] show the rather unusual and highly complex structure of 5S rDNA in *M. acuminata* using graph-based clusters of reads with multiple IGS and retroelement components, supporting the complexity reported here in the *E. glaucum* assembly. The multiple hybridization sites evident from the *in situ* hybridization site on one pair of chromosomes (Fig. 7B,C), with several, non-continuous, signals visible in the extended prometaphase chromosomes (Fig. 7C, insets) support the non-continuous nature of the 5S rDNA array.

The 5S rDNA monomer length of 1056bp was exceptionally long in comparison to any other plant species (typically 400-500 bp long). The first 400bp of the intergenic spacer had no significant BLAST hits in GenBank, while the second part showed only short regions with weak homology largely to chromosome assemblies of *Musa* species in GenBank. There were no motifs characteristic of retroelements in the 937bp intergenic spacer. It is unclear why the monomer length for the 5S rDNA in *E. glaucum* should be twice that typical in other species, including *Musa*, and was to be relatively homogeneous over all copies (Fig. 7E). Furthermore, in contrast to the single locus on eg04 of *E. glaucum*, all species of *Musa* examined so far have multiple 5S rDNA sites (2, 3 or 4 per genome), and *E. gilletii* had 3 pairs of sites [21, 47].

Different species in the Triticeae show wide variation in numbers and locations of both 45S and 5S rDNA sites, suggesting multiple and complex evolutionary rearrangements of the chromosome arms [50] even in the absence of chromosomal rearrangements including translocations and inversions. Dubcovsky and Dvorák [51] have considered the 45S rDNA loci as the “nomads of the Triticeae genomes” given their repeated evolutionary changes in position during species radiation without rearrangements of the genes of the linkage groups. The depletion of genes in chromosomal regions extending over most of a chromosome arm around the 45S rDNA genes is notable in *Musa* and *Ensete*, so chromosome rearrangements can lead to loci moving, although other recombination, duplication, deletion or translocation events must occur to alter the numbers of both 5S and 45S loci observed.

### Synteny and chromosome rearrangements to *Musa*

Structural comparisons of the *E. glaucum* genome sequence (x = 9, chromosomes eg01 to eg09) were performed with the high-quality assembled genomes of *M. acuminata* (x = 11, ma01 to ma11; v4, [20]) using translated proteins. SynVisio [52] was used for syntenic block visualization (Fig. 8A) and the comparison was extended to *M.* balbisiana (mb01 to mb11;[15]; Supplementary Fig. S8). Sequence dotplots (Fig. 8B) and comparative karyotypes (Fig. 8C and Supplementary Table S15) of the *E. glaucum* genome against the *M. acuminata* genome were also analyzed.

Overall, the genome assemblies of *M. acuminata* and *E. glaucum* are very similar in length and gene content (Supplementary Table S5). We observed high identity between segments of the 9 chromosomes of *E. glaucum* and of the 11 chromosomes of the *Musa* (Fig 8A). Broad centromeric regions with few protein-coding genomes (Fig. 2) cannot show syntenic domains. The number of collinear genes was 48,956 between *E. glaucum* and *M. acuminata,* and 39,604 between *E. glaucum* and *M. balbisiana* (by comparison, the A and B genome of *Musa* show 42,854 collinear genes). Chromosomes show rearrangements, inversions, expansions or contractions by crossed, converging or spreading lines in the Synvisio plots. The dotplot (Fig. 8B; single chromosome comparison in Supplementary Fig. S9) shows that there are some syntenic regions where the genes are distributed over the same length of chromosomes in both species (diagonal lines showing synteny at 45°, eg08/ma10), while in other cases there is expansion in one genome and not in the other (lines of synteny more vertical, eg04/ma04, or nearer horizontal, eg03/ma04). Many syntenic segments showed curved lines (ma03/eg03), showing relative expansion of one genome at one end of the conserved syntenic block, and expansion of the other genome at the other end.

One complete chromosome, ma05/eg05/mb05, was similar with the same gene content in all three species (Fig. 8A and Supplementary Fig. S8), but it showed multiple internal inversions and expansions/contractions. A nested pair of inversions was evident covering 10.4Mb near the start of the chromosome in *E. glaucum* with respect to *M. acuminata* (8.84Mb region) in dotplots (Supplementary Fig. S9A) and by comparison of locations of orthologous genes (Supplementary Fig. S10). In the context of the *E. glaucum* inversions, we could also examine the ancestral structure of *M. acuminata* and *M. balbisiana* reported by Wang et al. [15]. Notably, a major rearrangement involving an inversion between *M. acuminata* ma05 and *M. balbisiana* mb05 [15] was the same inverted region as found one eg05, with an additional nested inversion of 3.1Mb in eg05 (Supplementary Figs S8, S10). Using positions of orthologous genes at the boundaries of syntenic regions, the inversion structure between chromosomes eg05, ma05 and mb05 was clear. Regardless of the ancestral condition, the result indicates closely similar inversion breakpoints were involved (at the ends of the segment, Supplementary Fig. S10) twice during evolution.

Apart from chromosome 5, three further whole chromosomes of *M. acuminata* are represented largely by a single, whole chromosome regions/arms of *E. glaucum,* with some rearrangements occurring within the chromosomes (Fig. 8): ma01 is mainly the second (right) arm of eg01; and ma02 is mainly the second (right) arm of eg02 (Fig. 8A); ma11 is entirely the first (left) arm of eg09 (see details in the dot blot of Supplementary Fig. S9B). The other arms of these three *E. glaucum* chromosomes (eg01, eg02 and eg09) and the remaining six chromosomes are related to blocks of the remaining seven *Musa* chromosomes. Four *Musa* chromosomes have translocated fusions of segments of two *Ensete* chromosomes. ma09 has the intercalary region of eg07, with an intercalary segment of eg06 inserted within the eg07 region. ma10 includes parts of three *Ensete* chromosomes, while ma04 has four segments from *Ensete* chromosomes (Figs 8A, C). The 45S rDNA on eg06 and ma10 are surrounded by syntenic regions but are both depleted in genes (Fig. 2). In contrast, the 5S rDNA sites are not surrounded by other orthologous genes (see above). The non-reciprocal translocation noted by Wang et al. [15] of a terminal segment between ma03 and mb01 lies within a larger syntenic block shared between ma03 and eg03. This suggests that the translocation occurred in the *M. balbisia*na lineage (Supplementary Fig. S8).

In several chromosomes of both *E. glaucum* and *M. acuminata*, the breakpoints occur in the centromeric regions (e.g., in eg02, eg03, eg07, eg08 and eg09; ma01, ma06, and ma10; Fig 8A). While some breakpoints occur at or adjacent to centromeres, the exact relationship of any breakpoint to the centromere and Egcen or Nanica sequences is diverse. Notably, eg03 is spanning the centromere of three *Musa* chromosomes, ma03, ma04 and ma08, and in other cases centromere regions are different despite surrounding synteny (e.g., eg07, ma07 and ma09). Telomeric or sub-telomeric regions are conserved between the two species in 7 of the 18 *E. glaucum* chromosome arms (e.g., eg01/ma01, eg06/ma07, eg08/ma10; the dotplot homology lines end in the corners of the chromosomes, Figs 8B and Supplementary Fig. S9B). In other chromosomes, telomeres in *E. glaucum* are in intercalary regions of *Musa* (see eg03/ma03 with inversion, eg08/ma06 in Fig 8B; and in detail eg08 and ma10 in Supplementary Fig. 9C). The homology of the whole chromosome ma11 and the left arm of eg09 (see above) indicates a fusion/fission event with loss/gain of centromere and telomere function, but they are also predicted for the other rearrangements discussed above.

Song et al. [10] review the data on the basic chromosome number of the Zingiberales, concluding that x = 11 is most reasonable original basic number, with x = 9 as a derived basic number. With chromosome numbers of x = 9, 10 and 11 predominant in Musaceae, this family is particularly suitable to explore the nature and locations of chromosomal fusions and fissions that are predicted to often occur in similar position during karyotype evolution (e.g. in wheat [53]). The availability of high-quality genome sequences, based on the ONT, Hi-C and in *Musa* BioNano and PacBio technologies, will allow the nature of breakpoints in chromosome fission events to be investigated at the sequence level between the *Musa* x=11 and *Ensete* x=9 species, as well as being able to shed light to centromere and telomere function.

### Conclusions

We provide a chromosome-scale assembly of *Ensete glaucum,* a sister genus to *Musa*. This assembly is valuable to infer *Musa* genome evolution, enabling comparison with putative last common ancestors of *M. acuminata* (A genome) and *M. balbisiana* (B genome) at protein and chromosomal levels. Most striking was the multiple rearrangements of chromosome structures between *E. glaucum* and the *Musa* A and B genomes, with only 4 of the 11 *M. acuminata* chromosomes (and only 3 of the 11 in *M. balbisiana*) showing synteny with only one or part of one *E. glaucum* chromosome. Despite new insight to chromosome evolution, the sampling still makes difficult to conclude between ascending or descending dysploidy, but it should be clarified once additional chromosome scale genomes are released in Musaceae (in particular the Callimusa section with n=7, 9 and 10) and in sister clades of the Zingiberales.

As well as the complex chromosome rearrangements, repetitive sequences differ extensively between the *Musa* and *Ensete* genera. There is a major tandem repeat at the centromeres of only the *Ensete* species, showing lack of conservation of this key structural element of chromosomes, although both genera have multiple copies of the Nanica retroelement in centromeric regions. *E. glaucum* has only one 5S rDNA locus, with an unusually long monomer of 1056bp.

The complete sequence provides an accurate reference for the genus for gene identification, marker development, Genotyping-By-Sequencing and Genome Wide Association Studies (GWAS) and will accelerate our understanding of the molecular bases of traits such as cold tolerance and starch accumulation, and allow identification of relevant genes. The work builds towards a complete pangenome of the Musaceae family, defining structural, gene and genetic diversity which can be used for genetic improvement across the Musaceae and potentially more widely.

## Material, Methods and Validation

### Sample collection and distribution

The individual *Ensete glaucum* plant used for genome sequencing and analysis was collected from Puer city, Yunnan province, China and maintained in the South China Botanical Garden, Guangdong province, China (accession no. 19990288; Fig. 1A-E). The distribution of *Ensete* and *Musa* species were identified in databases of Flora of China, South China Botanical Garden, iNaturalist, GBIF [54] (excluding cultivation sites) and regional distributions maps. Fig. 1F was then made from POWO [55] (overlaid and color-adjusted in Adobe Photoshop CC2018).

### DNA extraction and sequencing

Young leaves of *Ensete glaucum* were collected and ground into powder in liquid nitrogen. High molecular weight genomic DNA was extracted using the DNeasy Plant Mini Kit

(Qiagen, Hilden, Germany). DNA quality was assessed by agarose gel electrophoresis and NanoDrop 2000c spectrophotometry, followed by Thermo Fisher Scientific Qubit fluorometry.

#### Illumina sequencing

A genomic DNA library with 400bp fragments was constructed using Truseq Nano DNA HT Sample preparation Kit (Illumina USA Preparation Kit (Illumina, USA), and 150bp paired ends were sequenced with Illumina Novaseq by Grandomics Research Services (previously known as Nextomics, Wuhan, PR China). After applying Trimmomatic v0.36 [56] to trim adaptors, filtering out low quality reads and further quality control with fastQC v0.11.9 [57], 246 million paired reads and 36.88Gb of data resulted (Table 1).

#### Oxford Nanopore (ONT) sequencing

ONT (Oxford Nanopore, Oxford, UK) sequencing was performed by Grandomics Biosciences Co., Ltd. (Wuhan, China): long fragments longer than 12 kb were selected with BluePippin (Sage Sciences), and the SQK-LSK109 kit (Oxford Nanopore) was used to build a library that was sequenced using PromethION. The base calling was performed with Guppy and reads mean_q score_template (Phred) > 7 (base call accuracy >80%) were selected. A total of 129 Gb ONT reads (∼ 250X coverage) was generated. fastp v0.19.7 [58] was used for quality control including adaptor-trimming, filtering reads with too many Ns or mean q score lower than 7 and resulted in remaining clean data of 109 Gb (Table 1). The mean read length was c. 20kb, with the longest >120kb (Supplementary Fig. S11).

#### Hi-C chromatin interaction data

The Hi-C library was prepared followed by a procedure with an improved modification [59]. Nuclear DNA from fresh leaves was first fixed and cross-linked in situ with formaldehyde [60], extracted, and digested by the restriction enzyme *Dpn*II. The resulting sticky ends were biotinylated, diluted, and ligated. The biotinylated fragments which contained DNA of two physically interacting regions were used to make a paired-end library and sequenced with Illumina HiSeq platform (performed by Grandomix, loc. cit.). Quality control of Hi-C raw data was performed using Hi-C-Pro v2.8.1 (HiC-Pro, RRID:SCR_017643): low-quality sequences (quality scores<20), adaptor sequences, and sequences shorter than 30 bp were filtered out using fastp v0.19.7 [58].

### Genome and chromosome assembly

ONT data were corrected by Nextdenovo v2.0-beta.1 (Nextomics, 2019) and the 109 Gb filtered data were assembled by SMARTdenovo (SMARTdenovo, RRID:SCR_017622) [61]. To polish the assembly, contigs were refined with Racon (Racon, RRID:SCR_017642) [62], BWA v0.7.17 (BWA, RRID:SCR_010910) [63] was used to map the filtered Oxford Nanopore reads to the assembly and NextPolish v1.3.1 [64] was used to discard possibly redundant contigs and generate a final assembly; similarity searches were performed with the parameters “identity 0.8 – overlap 0.8”. Finally, BWA and Pilon v1.21 [65] were used to further correct the assembly using the Illumina Novaseq reads, and two rounds of mapping back to the assembly each time with further correction were undertaken. A 494Mb assembly with 124 contigs was achieved (Table 2 and Supplementary Tables S1, S2).

Read pairs from the Hi-C data were mapped to the draft assembly using bowtie2 v2.3.2 (bowtie2; RRID:SCR_016368) [66] with the settings -end-to-end, -very-sensitive and -L 30 to select unique mapped paired-end reads. Valid interaction paired-end reads were identified by HiC-Pro v2.8.1 (HiC-Pro, RRID:SCR_017643) [67] and retained for further analysis while invalid read pairs, including dangling-end, self-cycle, re-ligation, and dumped products were discarded. The scaffolds were further clustered, ordered, and oriented onto pseudo-chromosomes by LACHESIS (RRID:SCR_017644) [68],with parameters as follows: cluster_min_re_sites = 100, cluster_max_link_density = 2.5, cluster noninformative ratio = 1.4, order min n res in trunk = 60, order min n res in shreds=60. Finally, placement and orientation errors exhibiting obvious discrete chromatin interaction patterns were manually adjusted. A Hi-C contact matrix plot (Supplementary Fig. S12) showed no discontinuities in the assembly.

Validation of the assembly was performed using Benchmarking Universal Single-Copy Orthologs with (BUSCO, RRID:SCR_015008) v5 [18] to assess the completeness and presence of 1614 genes in the embryophyta_odb10 database in “genome” mode (Supplementary Table S3a).

### RNA extraction, sequencing and transcriptome assembly

Total RNA was extracted from fresh leaves and roots of the same individual of *Ensete glaucum* that was used for genomic sequencing using RNeasy Plant Mini Kit (Qiagen). Illumina libraries were built from 1 µg total RNA of each sample with TruSeq RNA Library Preparation Kit (Illumina, USA) and were then sequenced using Illumina Novaseq platform to generate paired-end reads. A transcriptome assembly was produced using RNAseq data using Trinity (Trinity, RRID:SCR_013048) [69] and mapped on the genome with PASA (PASA, RRID:SCR_014656) [70].

### Genome size estimation

Using the Illumina DNA sequence, genome size was estimated from the 17-mer frequency using Jellyfish v2.0 (Jellyfish, RRID:SCR_005491) [71] with the formula k-num/k-depth (where k-num is the total number of 17-mers, 30,417,960,841; and k-depth the highest k-mer depth, 54; Table 2). 21-mer data were used in findGSE [72] and Genomescope 2.0 (Genomescope R, RID:SCR_017014) [73] to estimate the genome size and heterozygosity (Supplementary Fig. S1).

### Gene annotation

We adopted a combination of ab initio gene prediction, homology-based gene prediction and transcriptome-based gene prediction strategy. RepeatMasker v4.0.9 (RepeatMasker, RRID:SCR_012954) with option “-no_is -xsmall” was used to generate a repeat softmasked genome file. RNAseq data from leaf and root tissues were mapped to the masked genome assembly with STAR v2.7 (STAR, RRID:SCR_004463) [74]. The RNA alignment was input into BRAKER2 v2.1.5 (BRAKER, RRID:SCR_018964) [75], a combination of GeneMark (GENEMARK, RRID:SCR_011930) [76] and AUGUSTUS (RRID:SCR_008417) [77], to perform ab initio gene predictions. The gene models from BRAKER2 were inputted into MAKER v2.31.10 (MAKER, RRID:SCR_005309) [78] as model, and the RNA alignment of *E. glaucum* and proteins form *M. acuminata* v2 were used as EST and protein evidence, respectively. We also utilized GeMoMa v2.3 (GeMoMa, RRID:SCR_017646) [79] to perform homology-based gene prediction using Musa acuminata v2 [19] as reference annotated genome. EvidenceModeler (EvidenceModeler, RRID:SCR_014659) [80] was used to combine de novo and homology-based predictions and our transcriptome evidence to produce the final structural gene annotation.

To annotate the function of predicted genes, we performed BLASTP (e-value = 1e−10) (BLASTP, RRID:SCR_001010) from the BLAST+ package [81] for each predicted coding sequence against the databases: UniProtKB/Swiss-Prot, UniProtKB/TrEMBL [82] and NR (non-redundant protein database at NCBI). These sequences are then processed to produce a non-identical (often referred to as pseudo non-redundant) prediction. To assign a putative function to a polypeptide we kept only the best hit based on three parameters: (1) Qcov (Query coverage = length high-scoring segment pair (HSP)/length query), (2) Scov (Subject coverage = length HSP/length subject) and (3) identity. Additional functional information was added by scanning sequences with InterProScan v5.46 (InterProScan, RRID:SCR_005829) [83]. Blast2GO v6.0.1 (Blast2GO, RRID:SCR_005828) [84] was used to integrate the results of blast and InterProScan, and to link the GO (Gene ontology) terms to genes accordingly (Supplementary Table S4) The functional annotation procedure is given in greater detail at [85].

BUSCO (BUSCO, RRID:SCR_015008; v5.0.0) was run in mode “transcriptome” using the embryophyta_odb10 database to assess the gene annotation results and found 1529 (94.7%) complete BUSCOs (Supplementary Table S3b).

### Gene family analyses

#### Orthogroups (OGs) identification in Musaceae

Protein-coding genes from *M. acuminata* [19], *M. balbisiana* v1.1 [15] and *M. schizocarpa v*1 [13] were retrieved from the Banana Genome Hub [86]. Protein-coding genes predicted from *E. ventricosum* was downloaded at NCBI Genome (GCA_000818735.3) to allow discrimination of *Ensete* specific OGs *E. glaucum* specific OGs. Combined with *E. glaucum* protein-coding genes, we used OrthoFinder v2.5.2 (RRID:SCR_017118) [87] and Diamond [88] with default parameters (summary in Supplementary Fig. S5). Visualization (Fig. 3B) was produced with UpsetR [89]. Gene ontology (GO) enrichments were calculated using TopGO [90] with Fisher’s exact test (Supplementary Fig. S2 and Supplementary Table S8).

#### Gene family expansion and contraction

To identify gene family expansion and contraction, we expanded previous analyses with OrthoFinder by adding a representative of Musaceae sister clades in Palms (*Phoenix dactylifera,* date palm, [91]) and Poales (*Oryza sativa v7*, rice, [92]; data downloaded from Phytozome [93]) but omitting *E. ventricosum* due to gene redundancy. The longest transcripts were kept if alternative splicing occurred. Divergence time estimation with Approximate Likelihood Calculation used MCMCTREE in PAML v4.9j (PAML, RRID:SCR_014932). Computational Analysis of gene Family Evolution (CAFE v4.2.1, RRID:SCR_018924) [94] was used to model the evolution of gene family sizes and stochastic birth and death processes and summarized in the phylogenetic tree (Fig. 3C).

#### Transcription factors

Protein coding gene sequences for *E. glaucum* and *M. acuminata* v2 were searched in PlantTFDB v5.0 (PLANTTFDB, RRID:SCR_003362) and iTAK online v1.6 [95]. Predicted transcription factors (TFs) were verified through a Hidden Markov Model (HMM) with PFAM searching tools using the cutoff E-value of 0.01 (Fig. 3D and Supplementary Table S9). Genes were verified by PFAM (Pfam, RRID:SCR_004726), CDD (Conserved Domain Database, RRID:SCR_002077, and SMART (SMART, RRID:SCR_005026) databases.

### Whole-genome duplication (WGD)

To identify the whole-genome duplication events, we applied WGDI pipeline (whole-genome duplication identification v0.4.7 [96]). The predicted proteins of *E*. *glaucum* were blasted against themselves and then a collinearity analysis was conducted. The *K*s (the synonymous rates of substitution) between genes in paired collinearity gene groups were calculated and the *K*s peak was detected. For comparison, the same processes were also applied to *Musa acuminata* v2 [19].

### Synteny analyses

Structural comparisons of the *E. glaucum* genome were performed with *M. acuminata* v4 (designated the A genome) and *M. balbisiana* (B genome). The very recent release of *M. acuminata* v4 assembly was preferred for this analysis as it improved pericentromeric regions and provided telomere-to-telomere gapless chromosomes [20]. Assemblies were aligned with minimap2 (Minimap2, RRID:SCR_018550) [97] and visualized results using D-Genies (D-GENIES, RRID:SCR_018967) v1.2.0 [98]. Protein-coding genes were processed to identify reciprocal best hits (RBH) with BLASTP (e-value 1e-10) followed by MCScanX (e-value 1e-05, max gaps 25) [99] and results imported in SynVisio [52] for syntenic block visualization. Scale bars and coloring of the chromosome bars was adjusted using Adobe Photoshop CC2018.

The karyotype of *E. glaucum* in Fig. 8C, was prepared from lengths of each pseudo chromosome (Supplementary Table S2) with the estimated centromere position (Supplementary Table S13) to estimate the left (darker colored) and right (lighter colored) chromosome arms. Chromosome lengths and centromere positions for *M. acuminata* were taken from [20] (Fig. 2a); syntenic blocks were calculated using the SynVisio diagram (Fig. 8A).

### Repetitive DNA identification and annotation

For repetitive DNA analysis, publicly available programs (see below and [36]) as well as manual searches and sequence comparisons were applied. Geneious v.10.2.6 (Geneious, RRID:SCR_010519) (Biomatters Ltd., Auckland, New Zealand) was used to produce the dotplots of Figs 5A, B, 7E and 8B.

In the assembly, repeated sequences were first searched with REPET v2.5 pipeline [100]. The top 100 repeated sequences were plotted on the four reference *Musa* genome assemblies (i.e. *M. acuminata*, *M. balbisiana, M. schizocarpa* and *E. glaucum*) using BlastAndDrawDensity.py script described in [20] and available on the Github repository [101],

#### Transposable Elements

Transposable elements were annotated by EDTA pipeline [102], which integrates various software to discovers TE including: long terminal repeat (LTR) retrotransposons [103–105], terminal inverted repeat (TIR) transposons [106], short TIR transposons or miniature inverted transposable elements (MITEs) [107], and Helitrons [108]. According to suggestions in [109], we also adopt RepeatModeler2 (RepeatModeler, RRID:SCR_015027) v2.0.1 [109] to find remaining TEs.

The TE-denovo procedure was used on masked assembly to produce a batch of 4229 TE consensus sequences. From these 2800 consensus sequences, only those with full length fragments present in the assembly were kept for further analysis, quantification and annotation with the TE-ANNOT procedure. A first annotation was performed using public Repbase (Repbase, RRID:SCR_021169) release 20.05, followed by *Gypsy/Copia* retroelement family identification using Hidden Markov Models (hmmsearch version 3) to search consensus for corresponding retrotransposase PFAM domains PF04195 and PF14244 respectively. The above results were then combined and CD-HIT v4.1.8 (CD-HIT, RRID:SCR_007105) [110] was used to reduce redundancy. The LTR retrotransposons were sent to TEsorter [111] to classify into lineage level and RT domain amino acid sequences were extracted. Phylogenetic trees of *Copia* and *Gypsy* were inferred by RT domain alignment results (Supplementary Figure S4). The proportions of transposons in the assembly are given in Table 2 (Supplementary Tables S11 and S12) and chromosomal distributions in Fig. 2.

To estimate ages of LTR transposons and the time of insertion (Fig 4B, C and Supplementary Figure S5), complete elements were found by LTRharvest v1.6.1 (LTRharvest, RRID:SCR_018970) [103] and LTR_retriever (LTR_retriever, RRID:SCR_017623) [104] and then classified by TEsorter v1.3. The estimation of time was based on the divergence of of the 5’ and 3’ end LTRs, and these two LTRs of every LTR retrotransposons were extracted into separate files with a custom script. The 5’ and 3’ LTRs were aligned by MUSCLE v3.8.1551 [112]. The divergence distances under K2P evolutionary model were calculated by R package ape v5.4-1 (ape, RRID:SCR_017343). The average base substitution rate was selected to be 11.3E-8 [113]. The insertion time T was calculated as T = K/(2r), with r as the rate of nucleotide substitution and K as the divergence distance between LTR pairs. Our script to perform the analysis is on github (2021) [114].

#### Graph based clustering of reads using RepeatExplorer

A sample of 2Gb of the Illumina HiSeq raw reads were used for assembly-free analysis by RepeatExplorer2 [37]. Graph-based clusters of similar sequence fragments were generated under default parameters. Clusters were assigned to repeat classes and retroelement lineages using the automated Repeat Masker and Domain hits provided by the program (Supplementary Table S11). Comparative analysis with sample Illumina sequence reads from other five Musaceae species, namely *M. acuminata* v2 [19], *M. balbisiana* v1.1 [15] and *M. schizocarpa* v1 [13], *E. ventricosum* [17] and *Musella lasiocarpa* (Ziwei Wang, Qing Liu and Pat Heslop-Harrison pers communication) were also analyzed (Supplementary Figure S3) and compared with RepeatExplorer2 following “comparative repeat analysis” protocol. The results were visualized by R script “plot_comparative_clustering_summary.R” (Fig, 4A).

#### SSR Tandem Repeats

The genome assembly was searched for SSR (microsatellite) motifs using the SSR mining pipeline developed by Biswas et al [46]. Searches were standardized for mining perfect SSRs from mono to hexa-nucleotide repeats (minimum repeat number of 12 for mononucleotides, 8 for di-, 5 for tri-, tetra- and penta-, and 4 repeats for penta- and hexa-nucleotides). SSRs abundance and nature was analyzed based on density in the genome (about 1 per 4000bp), array length (Figure 7B and Supplementary Table S14; SSR search parameters minimum lengths mono=1*12=12nt, di=8*2=16nt, tri=3*5=15nt, tetra=4*5=15nt, penta=5*4=20nt and hexa=6*4=24nt; total SSR count 123884; Class I>20nt and Class II≤20nt), nucleotide base composition of the SSR loci (70% were AT-rich) and abundance of each motif).

### Fluorescent *in situ* hybridization (FISH)

Chromosome preparation and FISH was performed as described by Schwarzacher and Heslop-Harrison [115] with minor modifications. Plants of *E. glaucum, E. ventricosum, Musella lasiocarpa* (purchased commercially) and *M. balbisiana* ‘Butuhan’ (ITC1074) [116] were grown in the glasshouse at the University of Leicester, UK. Actively growing root tips were treated with 2 mM 8-hydroxyquinoline and fixed with 96% ethanol:glacial acetic acid (3:1). For chromosome preparations, roots were digested with a modified enzyme solution (32U/ml cellulose, Sigma-Aldrich C1184; 20U/ml ’Onozuka’ RS cellulose; 35U/ml pectinase from *Aspergillus niger,* Sigma-Aldrich P4716; 20U/ml Viscozyme, Sigma-Alderich V2010) in 10mM citric acid/sodium citrate buffer (pH4.6) for 3-5h at 37°C and then kept in buffer for 12-30h at 4°C. Meristems were dissected in 60% acetic acid and routinely 2-6 slide preparations were made from each root. Slides were stored at -20°C until FISH.

The 45S rDNA probe was labelled by random priming (Invitrogen) with digoxigenin dUTP (Roche) using the linearized clone pTa71 (Gerlach and Bedbrook 1979) containing the 45S rDNA repeat unit of *Triticum aestivum*. 50-100ng of labelled probe was used per slides and detection of hybridization sites was carried out with Fluorescein-conjugated anti-digoxigenin (Roche). The remaining probes were designed from the consensus sequence of the centromeric repeat Egcen (Fig. 5) and the 5S rDNA (Fig. 7) or as simple sequence repeats (Fig. 6); as directly labelled oligonucleotides (200-500ng per slide) they needed no further detection and were as follows:

CenCy3: EGL_2640R: [Cyanine3]GAC CGT CGC ATT TTT TGG CGA AAC CAT GCT CGT ACG ACT TCC CAT GGG CTA AAA CGT TAG GA

CenFAM: EGL_G2640L: [6FAM]GGC CTA TAT TTT GAA ATT CCG AGA CGG TGC ATG AAA AAC CGA TCG AAA CGA AAC ATT GCG

5S_4M_Cy3: [Cyanine3]TCA GAA CTC CGA AGT TAA GCG TGC TTG GGC GAG AGT AGT AC

5S_3R_Cy3: [Cyanine3]AGT ACT AGG ATG GGT GAC CCC CTG GGA AGT CCT CGT GTT GC

5S_6L_Cy3: [Cyanine3]GCG ATC ATA CCA GCA CTA AAG CAC CGG ATC CCA TCA GAA CTC C

(AAG)15_FAM: [6FAM]AAG AAG AAG AAG AAG AAG AAG AAG AAG AAG AAG AAG AAG AAG AAG

(CT)23_TAMRA: [TAMRA]CTC TCT CTC TCT CTC TCT CTC TCT CTC TCT CTC TCT CTC TCT CTC T

For hybridization, probes were prepared in 40% (v/v) formamide, 20% (w/v) dextran sulphate, 2x SSC (sodium chloride sodium citrate), 0.03 μg of salmon sperm DNA, 0.12% SDS (sodium dodecyl sulphate) and 0.12mM EDTA (ethylenediamine-tetra acetic acid). Chromosomes and 40-50µl of probe mixture were denatured together at 72oC for 8 mins, cooled down slowly and allowed to hybridize overnight at 37°C. Post-hybridization washes were at 42°C in 0.1xSSC, giving a stringency of 80-85% for the short oligo probes, and 70-75% for the 45SrDNA probe. Chromosomes were counterstained with 4µg/ml DAPI (4!,6-diamidino-2-phenylindole) and mounted in CitifluorAF. Slides were examined using Nikon Eclipse 80i microscope and images were captured with a DS-QiMc monochrome camera, and NIS-Elements v2.34 (Nikon, Tokyo, Japan). Overlays of hybridization signal and DAPI images were viewed enhanced with Adobe Photoshop CC2018 using only cropping and functions that treat all pixels equally. Seven FISH runs with different combinations of probes and replicates were performed, and between 5 and 15 metaphases per slide (99 metaphases in total from 15 slides) were analyzed in detail.

## Availability of Supporting Data and Materials

Raw sequence reads (RNA-seq, Illumina HiSeq, the Oxford Nanopore and Hi-C) were deposited in the NCBI under accession number: PRJNA736572. In specifically, the Oxford Nanopore raw reads: SRR15039768 and SRR15039769; RNA-sequencing raw reads, as follows: leaf: SRR15039767, root: SRR15039766; genomic Illumina short-read data: SRR15039770; raw reads of the Hi-C library: SRR15039764 and SRR15039765.The raw reads data were also deposited in Genome Sequence Archive (GSA) of the China National Center for Bioinformation (accession code: CRA004283).

The assembled genome was also deposited to GenBank in NCBI under the accession number: JAHSUZ000000000. Genome Assembly, gene and TE annotation data, transcriptomic data are also available on the Banana Genome Hub (http://banana-genome-hub.southgreen.fr/) for download or exploration via a dedicated Genome Browser (Jbrowse) and syntenic browser (SynVisio).

## Additional Files

### Supplementary Figures

#### Pdf -word document containing Supplementary Figures S1-S12

Supplementary Figure S1. Genomescope analysis of heterozygosity.

Supplementary Figure S2. GO enrichment terms.

Supplementary Figure S3. RepeatExplorer clustering summary in Musaceae species.

Supplementary Figure S4. LTR retroelement trees EGL and MAC.

Supplementary Figure S5. Gypsy and Copia insertion times in *Musa*.

Supplementary Figure S6: Egcen FISH to *Musa* chromosomes.

Supplementary Figure S7. EgCen and Nanica in Assemblies of *E. glaucum* and *Musa*.

Supplementary Figure S8 Synteny of *E. glaucum* with *Musa* A and B genome.

Supplementary Figure S9. Dotplots of individual chromosomes.

Supplementary Figure S10. Inversions on chromosome 5 in three Musaceae species.

Supplementary Figure S11. Length distribution of ONT reads.

Supplementary Figure S12. Hi-C interaction contact map.

### Supplementary Tables

#### Excel Spreadsheet containing Supplementary Tables S1-S14

Supplementary Table S1. Contig statistics based on assembly of ONT sequencing data.

Supplementary Table S2. Chromosome lengths and number of contigs anchored in *Ensete glaucum* assembly.

Supplementary Table S3. Quality assessment of the gene annotation of *Ensete glaucum* using BUSCOs v5.

Supplementary Table S4. Complete gene list. homology and GO.

Supplementary Table S5. Statistics for shared orthogroups (OG) and gene clustering among *E. glaucum, E. ventricosum, Musa acuminata, M. balbisiana,* and *M. schizocarpa* genomes.

Supplementary Table S6. Positively selected genes and their annotation.

Supplementary Table S7: Result of gene family size change analysis.

Supplementary Table S8 a) Top 20 GO molecular function enrichments for *E. glaucum* and shared *E. glaucum/E. ventricosum* gene families; b) Top 20 GO biological pathways enrichments for *E. glaucum* and shared *E. glaucum/E. ventricosum* gene families.

Supplementary Table S9. Comparison of transcriptional factor between *Ensete glaucum* and *Musa acuminata*.

Supplementary Table S10. Transposable elements and other repeat proportions comparison in assembly (RepeatMasker).

Supplementary Table S11. Repeat content (RepeatExplorer) comparison between different Musaceae genomes.

Supplementary Table S12. Abundance of major tandemly repeated DNA repeats in Illumina raw reads.

Supplementary Table S13. Inferred centromere positions from locations of interrupted tandem arrays of the Egcen centromeric sequence on the chromosome assemblies.

Supplementary Table S14. Comparative survey of microsatellite sequences in *Ensete glaucum* genome with other sister species.

## Declarations

## List of abbreviations

BLAST: Basic Local Alignment Search Tool
bp: base pairs
BUSCO: Benchmarking Universal Single-Copy Orthologs
BWA: Burrows-Wheeler Aligner
FISH: fluorescence in situ hybridization
GeMoMa: Gene Model Mapper
Gb: gigabase pairs
GC: guanine-cytosine
GO: gene ontogeny
CTAB: cetyl trimethylammonium bromide
GWAS: Genome Wide Association Studies
Hi-C: High-throughput chromosome conformation capture
ITS: internal transcribed spacer of rDNA
kb: kilobase pairs
KEGG: Kyoto Encyclopedia of Genes and Genomes
GeMoMa: Gene Model Mapper
LACHESIS: Ligating Adjacent Chromatin Enables Scaffolding In Situ
LINE: long interspersed nucleotide elements
LTR: long terminal repeat
Mb: megabase pairs
ML: maximum likelihood
miRNA: microRNA
Mya: million years ago
NCBI: National Center for Biotechnology Information
NR: RefSeq non-redundant proteins
NOR: Nucleolar Organizing Region
NTS: non transcribed spacer of rDNA
ONT: Oxford Nanopore Technologies
PAML: Phylogenetic Analysis by Maximum Likelihood
PacBio: Pacific Biosciences
PASA: Program to Assemble Spliced Alignments
RAxML: Randomized Accelerated Maximum Likelihood
RNA-seq: RNA sequencing
rDNA: ribosomal DNA
SRA: Sequence Read Archive
SSR: Simple sequence repeat
TE: transposable element
TF: Transcription Factor
tRNA: transfer RNA
WGD: whole genome duplication.

## Consent for publication

The origin of *E.glaucum* plants is given in Materials and Method. They were collected in China and conserved in the South China Botanical Garden, Chinese Academy of Sciences, with appropriate agreements. No live material was exported out of the country. Other plants for chromosome preparations was obtained from the International Transit Centre, ITC genebank, with official Standard Material Transfer Agreement (SMTA), acknowledged in the manuscript.

## Author contributions

Q.L. and J.S.H.H. designed the project and with M.R. and Z.W. contributed to project coordination. Z.W. and Q.H. collected samples and conducted DNA and RNA extraction. T.S. and J.S.H.H. conducted FISH experiments. Z.W. carried out genome assemblies; Z.W., M.R. G.D., and M.K.B. conducted gene annotation and comparative genomic analyses; Z.W., T.S., J.S..H.H. and F.C.B. conducted repetitive sequence analysis. All authors contributed to writing and editing the manuscript.

## Acknowledgments

This work was technically supported by the high-performance cluster of the UMR AGAP - CIRAD of the South Green Bioinformatics Platform (http://www.southgreen.fr). Assistance and discussion with Celia Hansen and Paulina Tomaszewska, University of Leicester, are acknowledged. We thank Ye Yushi for the assistance with growing and maintaining the plants. We thank the International Musa Germplasm Transit Centre (https://www.bioversityinternational.org/banana-genebank/) for samples of *M. balbisiana*.

