## Supplemental figures for "A chromosome-level reference genome of *Ensete glaucum* gives insight into diversity, chromosomal and repetitive sequence evolution in the Musaceae"

### S1. Genomescope analysis of heterozygosity

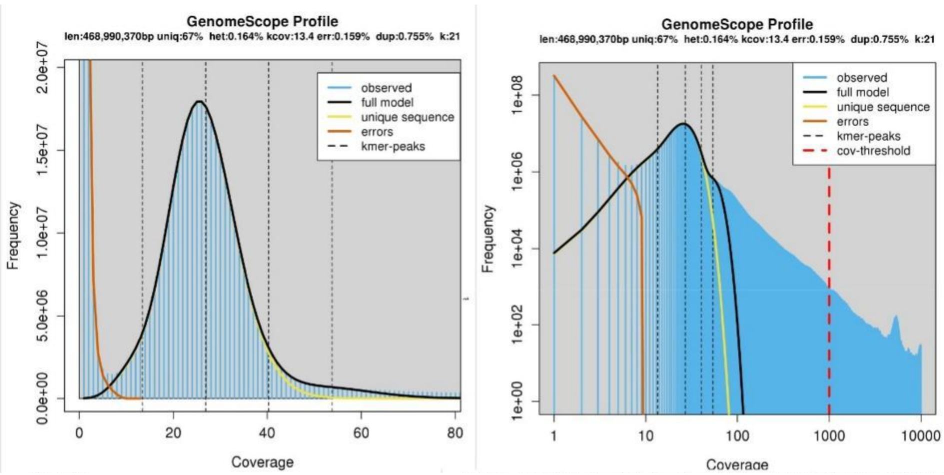

**Supplementary Figure S1: Heterozygosity and genome size estimated from Illumina raw-reads and 21-mer analysis using GenomeScope**. See genomescope for complete calculations.

### S2. GO enrichment terms

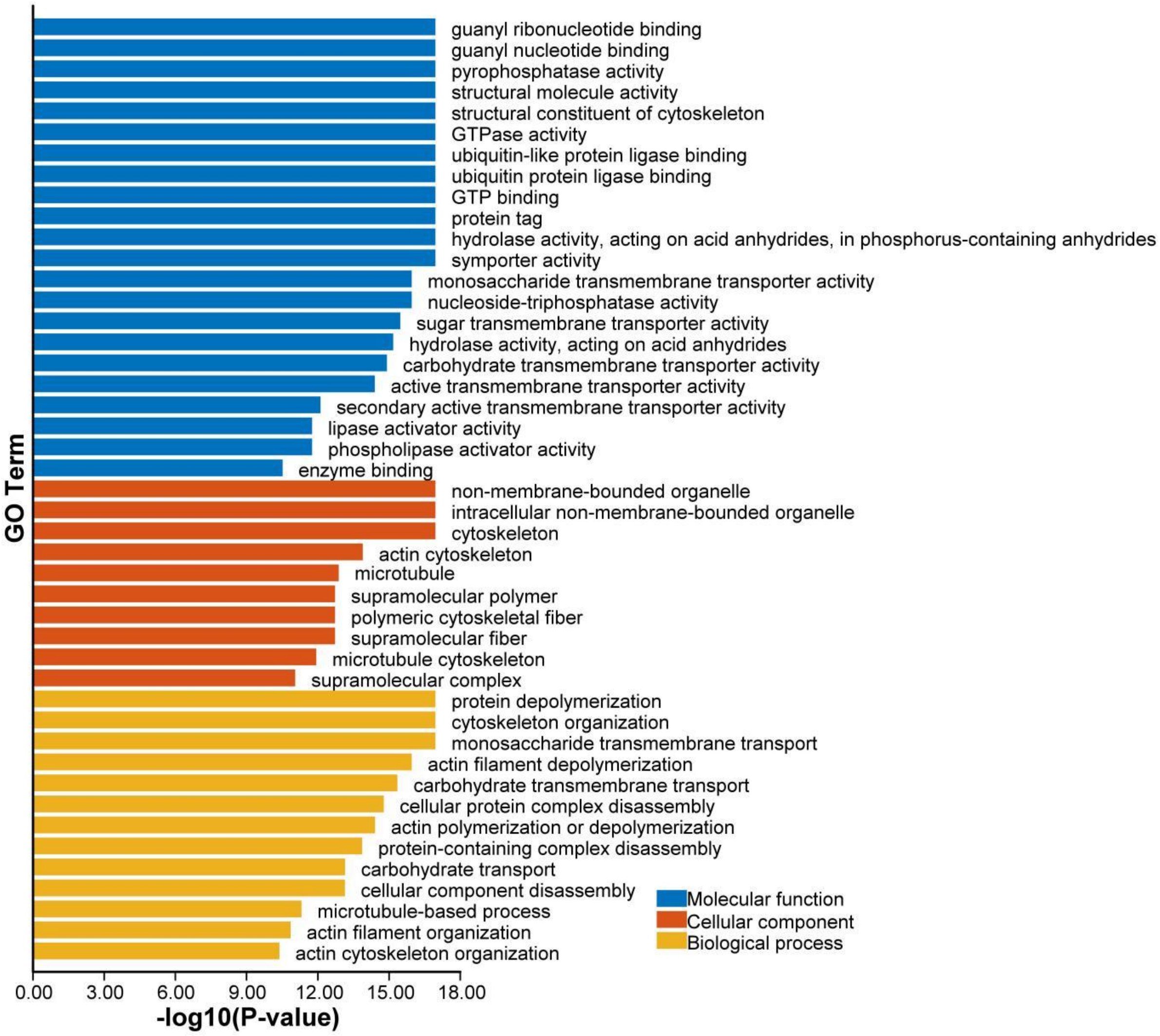

**Supplementary Figure S2: Top 50 GO enrichment terms of genes in rapidly expanding gene families in *E. glaucum*.**

### S3. RepeatExplorer clustering summary in Musaceae species

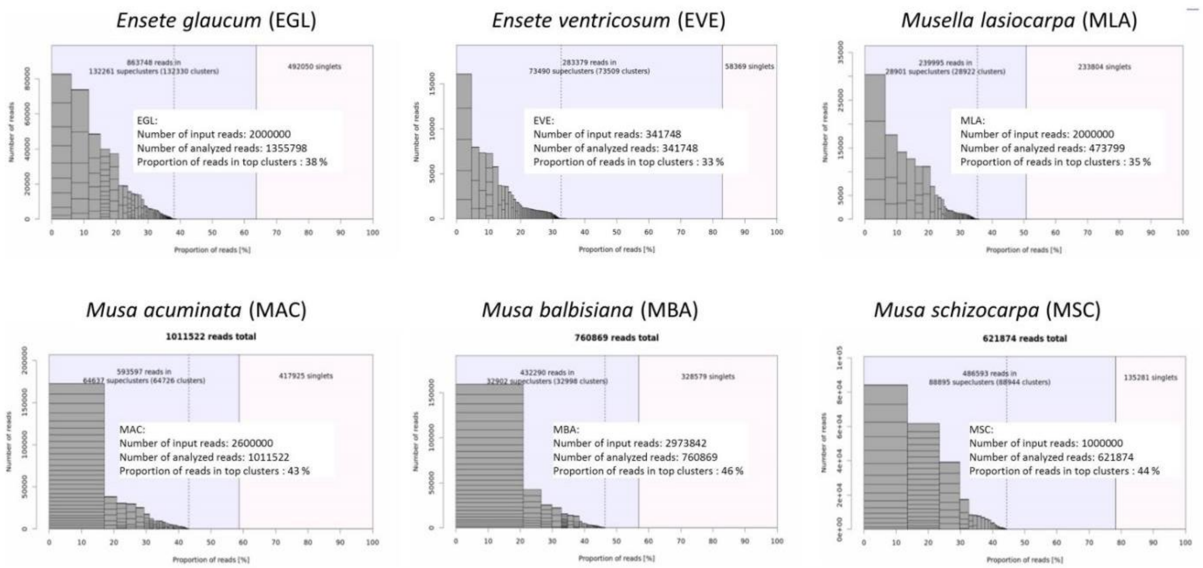

**Supplementary Figure S3: RepeatExplorer cluster analysis of raw reads of *E. glaucum*, *E. ventricosum*, *Musella lasiocarpa*, *M. acuminata*, *M.blabisiana* and *M schizocarpa*.** Summary of reads that are assigned to clusters or singleton (graphs), with numbers also given for clusters with genome frequency of > 0.01% (top clusters as defined by Novak et al. 2020).

### S4. LTR retroelement trees EGL and MAC

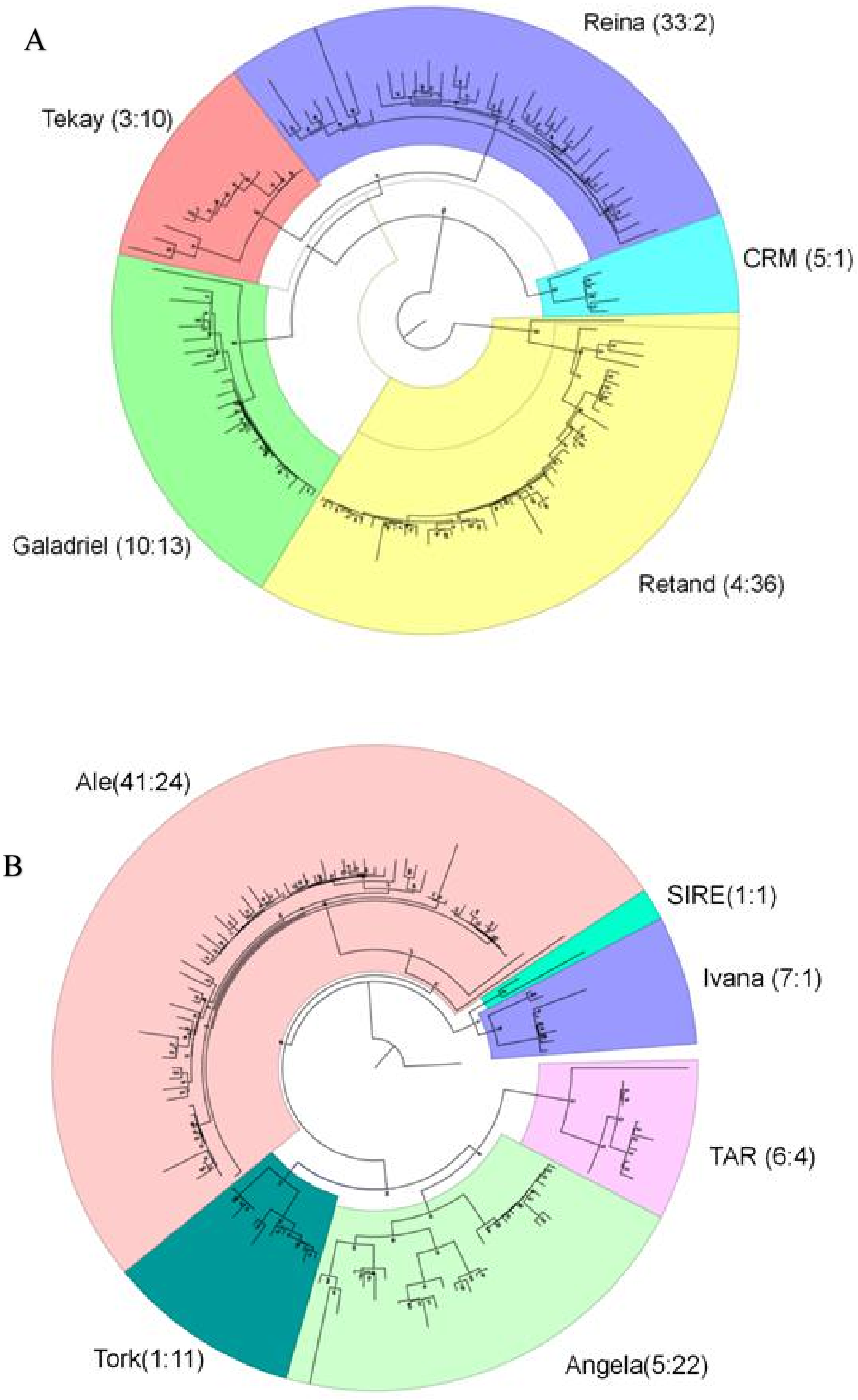

**Supplementary Figure S4: Phylogenetic tree of LTR-retroelements (A) Gypsy (B) Copia in *E. glaucum* and *M. acuminata* based on RT domains.** The number in brackets after each lineage represent the element numbers for *E. glaucum* (left) and *M. acuminata* (right)

### S5. *Gypsy* and *Copia* insertion times in *Musa*

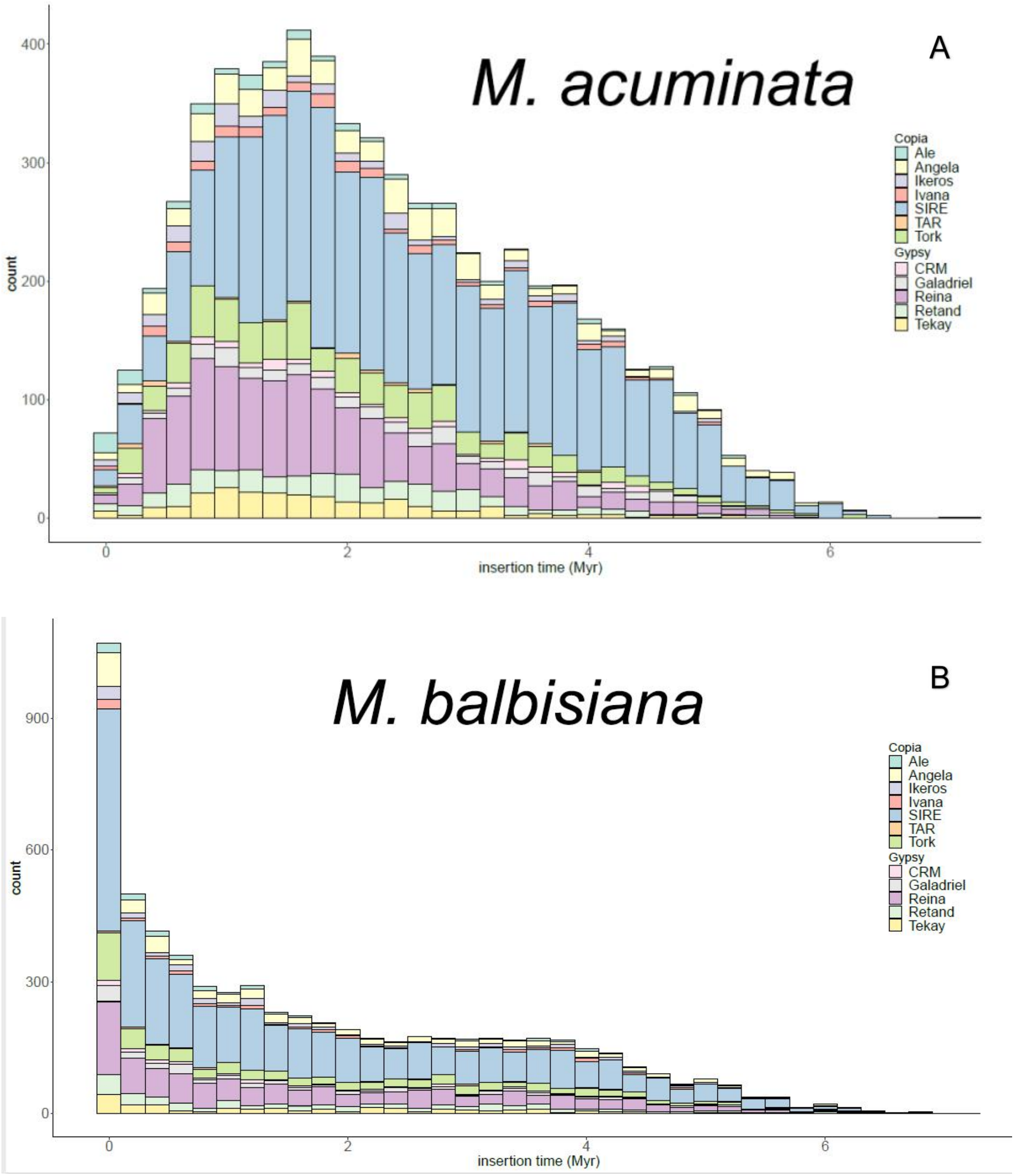

**Supplementary Figure S5: Insertion times of LTR-*Copia* and LTR-*Gypsy* retroelement calculated for (A) *M. acuminata* and (B) *M. balbisiana*.** For comparison with *E. glaucum*, see Fig.4B and C.

### S6: Egcen FISH to Musa chromosomes

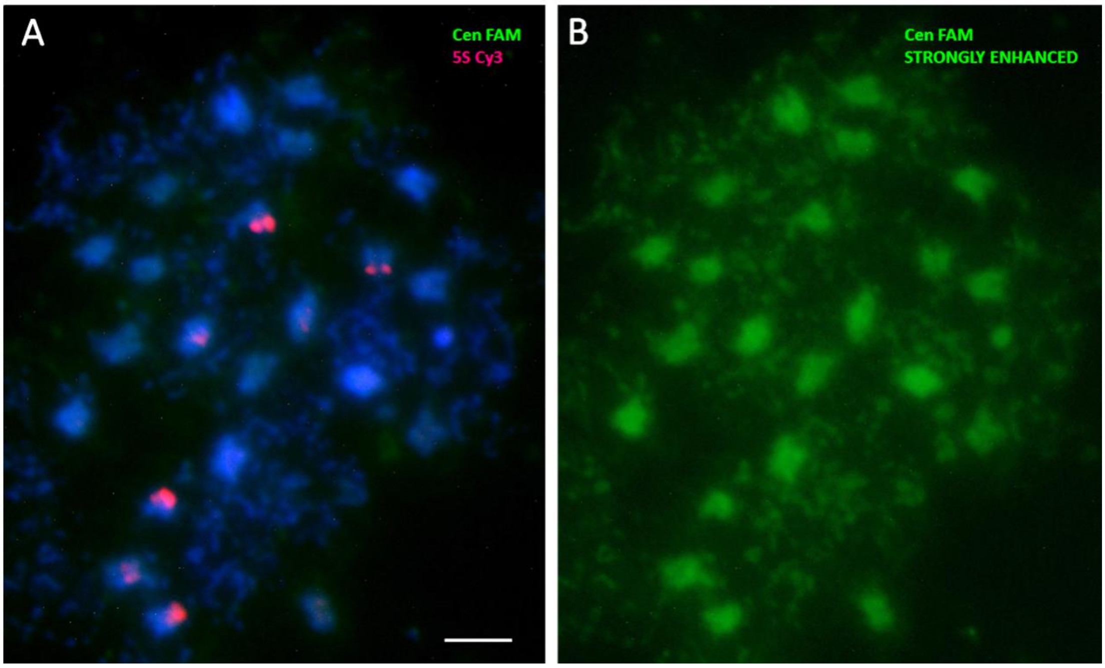

**Supplementary Figure S6: Metaphase chromosomes of *Musa balbisiana* 'Butuhan' 2*n*=22**
Fluorescent *in situ* hybridization using Egcen oligonucleotide labelled with FAM (green) and 5S oligonucleotide with Cy3 (magenta) and DAPI staining (blue). (A) Normal exposure and enhancement similar to images of *Ensete* and *Musella* (Fig. 6D-F) using the same probe combination processed in the same FISH run. (B) Strongly enhanced green image showing residual fluorescence that mimics the DAPI staining but does not show any real FISH signal of the Egcen probe. Bar 5 µm

### S7. Egcen and Nanica in assemblies of *E. glaucum* and *Musa*

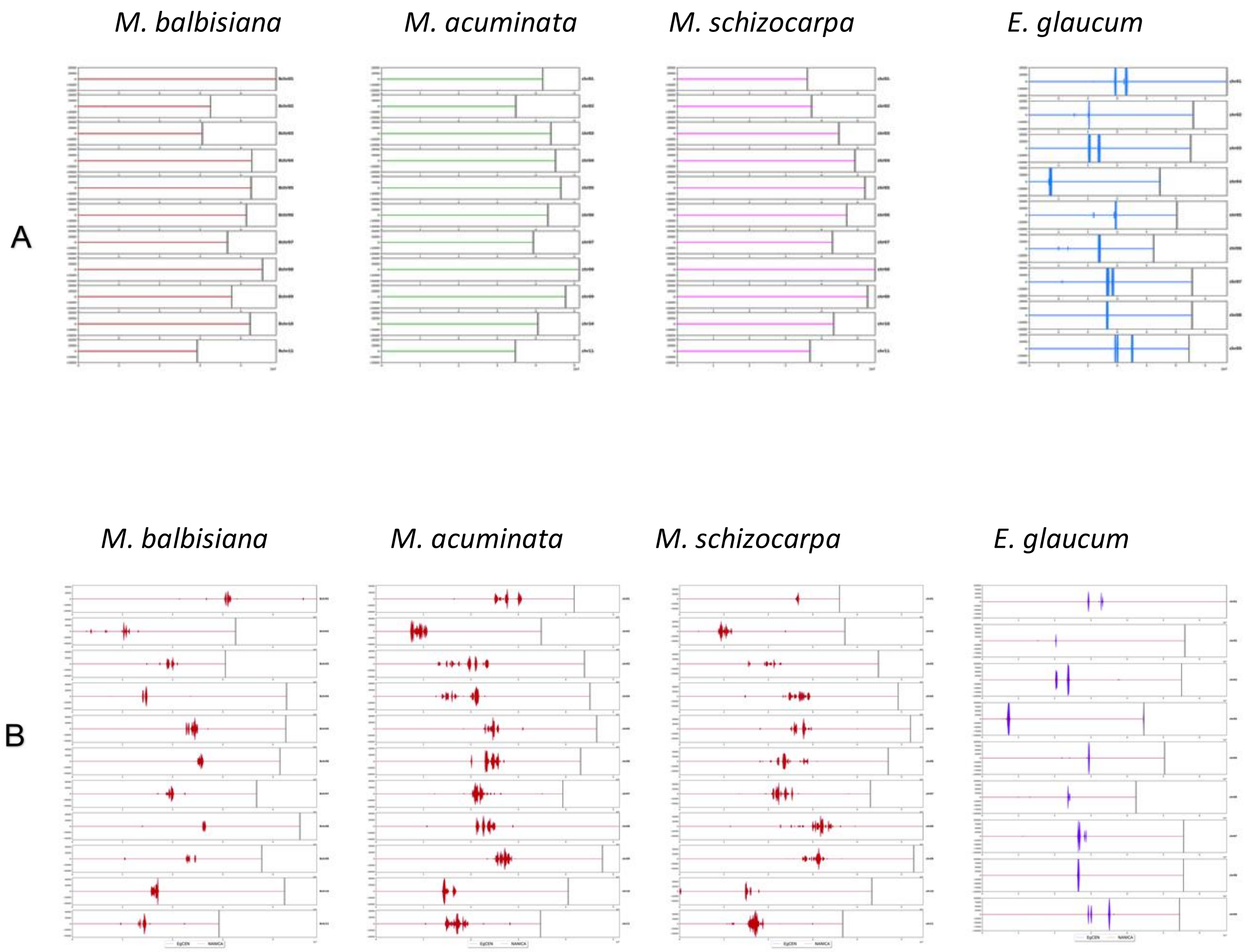

**Supplementary Figure S7: Graphic representation of Egcen and the LINE non-LTR retroelement Nanica in assemblies of three *Musa* genomes and *E. glaucum*.** (A) Egcen (vertical blue bars) has not been found in *Musa* assemblies (B) The non-LTR retroelement Nanika (vertical red bars) is found in all *Musa* species and with low copies also in *E. glaucum* at sites of Egcen (blue) compare with Fig. 5C.

### S8. Synteny of *E. glaucum* with *Musa* A and B genomes

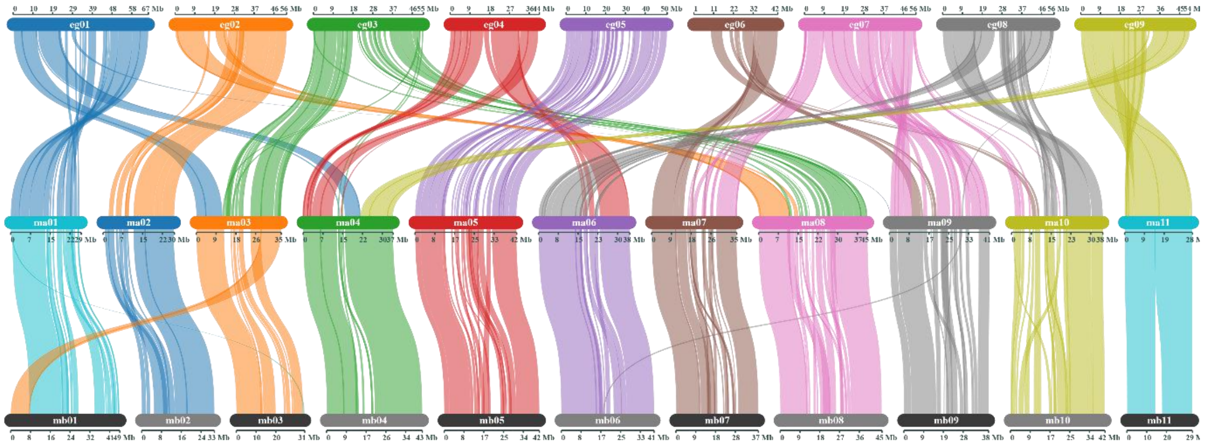

**Supplementary Figure S8: Synteny plot of the *E. glaucum* (top), *M. acuminata (v4;* Belser et al. 2021; middle) and *M. balbisiana (v1.1; Wang et al 2019;*(bottom)** Syntenic relationships were visualized by Synvisio. See also Fig. 8A.

### S9. Dotplot of individual chromosomes

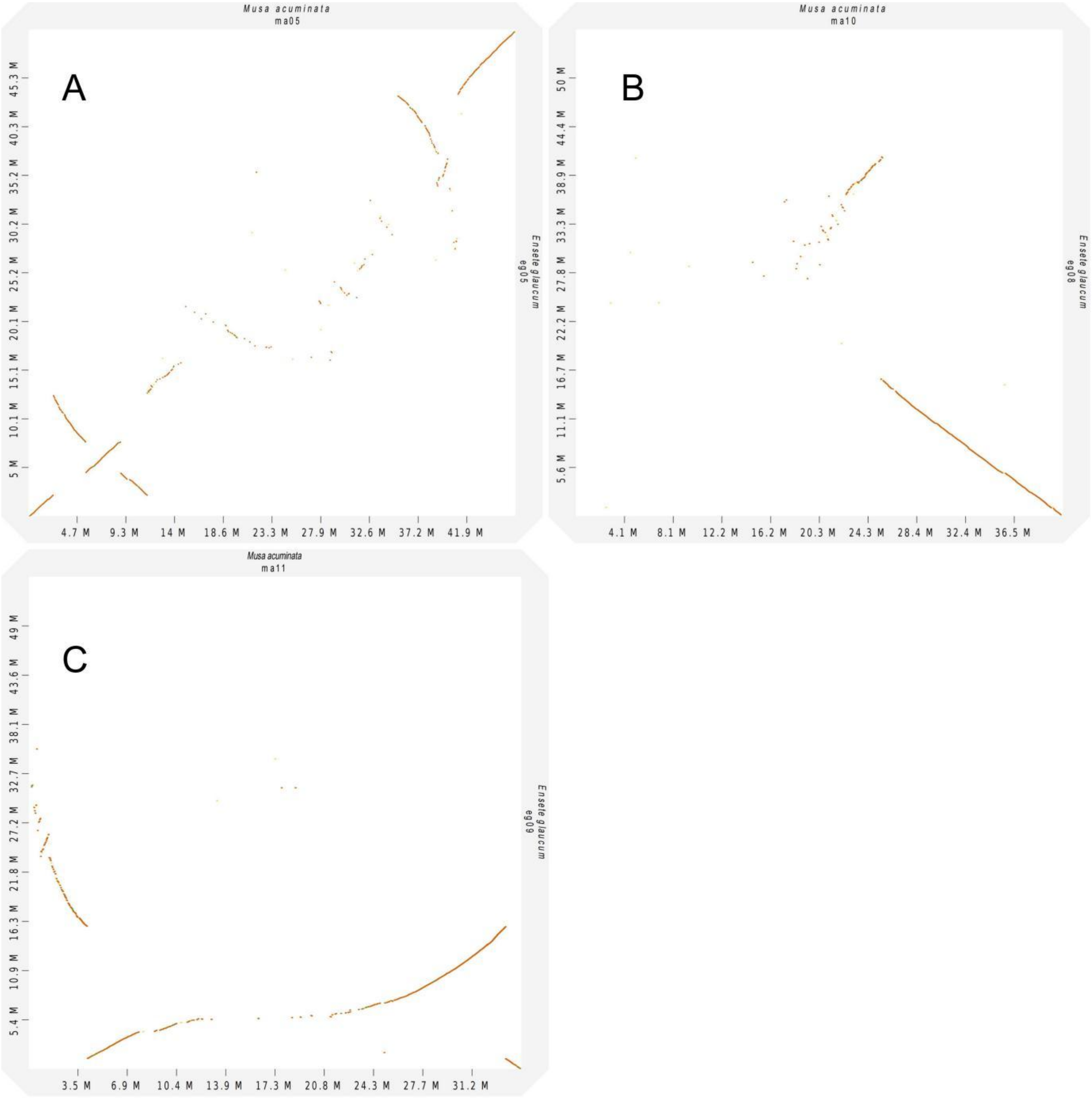

**Figure S9: Dotplots between individual chromosome assemblies of *E. glaucum* and *M. acuminata.*** Straight diagonal lines show high homologies. (A) eg05 and ma05 showing inversions and several small rearrangements (B) eg09 and ma11: the whole chromosome of ma11 covers one arm of eg09, but curved homolgy lines indicate amplification/deletion in one of the chromosomes. (C) eg08 and ma10. Matching telomeres are indicated by the line ending in the corner.

### S10. Inversions on chromosome 5

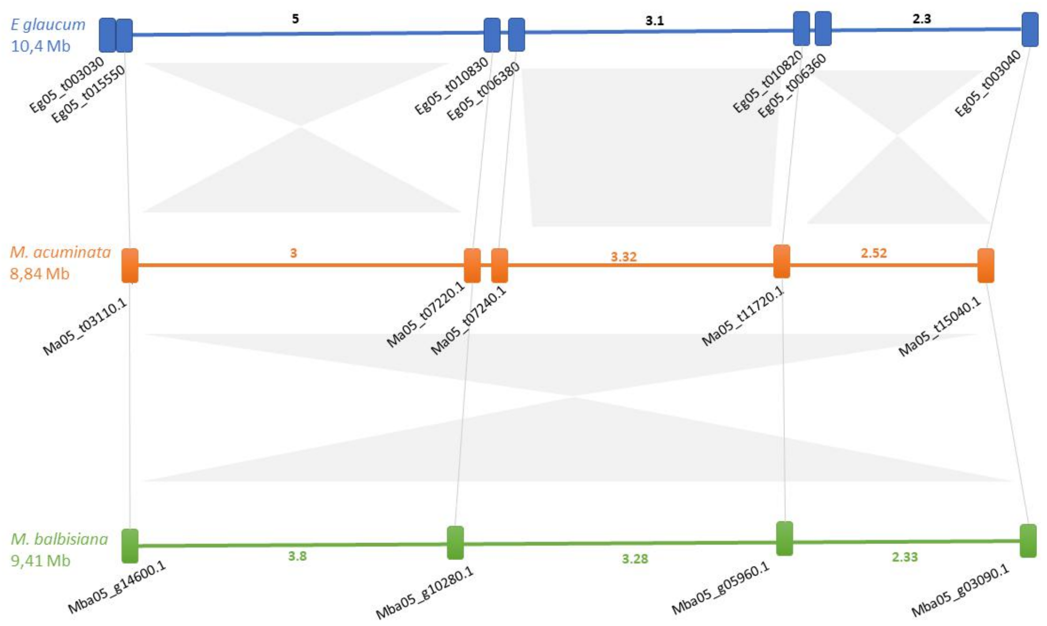

**Supplementary Figure S10: Schematic representation of chromosome 5 regions bearing inversions between *Musa acuminata, M. balbisiana* and *E. glaucum* showing nested inversion(s).** Syntenic genes for the three species at boundaries of are represented by rectangles with their locus tag linked by grey lines. Number on the segment indicates the length in Mb. Gray bars between species indicate conserved regions with filled areas showing synteny and symmetrical filled triangles symbolizing inversions.

### S11. Length distribution of ONT reads

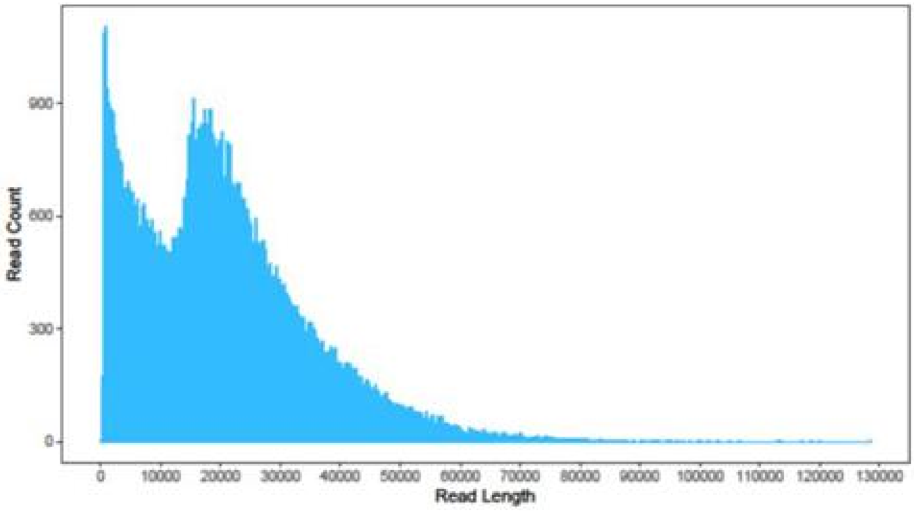

**Supplementary Figure S11: Distribution of ONT read length showing a peak at ~20,000 bp and the longest read length at ~130,000bp.**

### S12. Hi-C interaction contact map

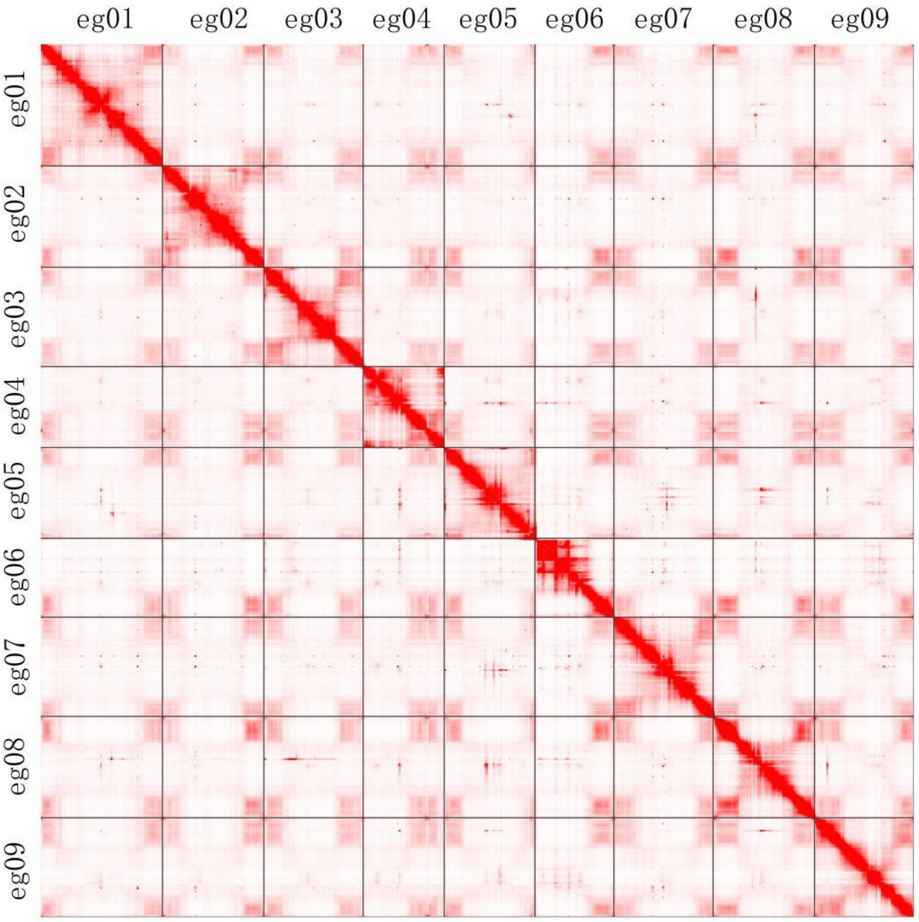

**Supplementary Figure S12: Hi-C interaction analysis depicting the 9 pseudochromosomes of *Ensete glaucum* genome.**
