## Supplemental tables for "A chromosome-level reference genome of *Ensete glaucum* gives insight into diversity, chromosomal and repetitive sequence evolution in the Musaceae"

**Supplementary Table S1. Contig statistics based on assembly of ONT sequencing data.**

| Stat Type | Contig [illegible] | Contig Number |
|---|---|---|
| N50 | 10,255,891 | 14 |
| N60 | 8,931,940 | 19 |
| N70 | 7,071,257 | 25 |
| N80 | 5,764,441 | 33 |
| N90 | 2,596,489 | 47 |
| Longest | 31,226,749 | |
| Total | 495,175,598 | 124 |
| Length>=1kb | 495,175,598 | 124 |
| Length>=2kb | 495,175,598 | 124 |
| Length>=5kb | 495,175,598 | 124 |

**Supplementary Table S2. Chromosome lengths and number of contigs anchored in Ensete glaucum assembly.**

| chromosome | chr.length/bp | contigs |
|---|---|---|
| eg01 | 67,484,389 | 11 |
| eg02 | 55,950,575 | 9 |
| eg03 | 55,030,165 | 7 |
| eg04 | 44,624,142 | 10 |
| eg05 | 50,342,496 | 13 |
| eg06 | 42,457,113 | 12 |
| eg07 | 55,595,191 | 12 |
| eg08 | 55,557,465 | 9 |
| eg09 | 54,465,677 | 7 |
| Total | 481,507,213 | 90 |

**Supplementary Table S3a. Quality assessment of the assembled genome of Ensete glaucum using BUSCOs v5 in 'genome' mode**

| Type | Number | Percent (%) |
|---|---|---|
| Complete BUSCOs (C) | 1587 | 98.3% |
| Complete and single-copy BUSCOs (S) | 1526 | 94.5% |
| Complete and duplicated BUSCOs (D) | 61 | 3.8% |
| Fragmented BUSCOs (F) | 12 | 0.7% |
| Missing BUSCOs (M) | 15 | 1.0% |
| Total BUSCO groups searched | 1614 | 100.0% |

**Supplementary Table S3b. Quality assessment of the gene annotation of Ensete glaucum using BUSCOs v5 in 'transcriptome' mode**

| Type | Number | Percent (%) |
|---|---|---|
| Complete BUSCOs (C) | 1529 | 94.7% |
| Complete and single-copy BUSCOs (S) | 1456 | 90.2% |
| Complete and duplicated BUSCOs (D) | 73 | 4.5% |
| Fragmented BUSCOs (F) | 48 | 3.0% |
| Missing BUSCOs (M) | 37 | 2.3% |
| Total BUSCO groups searched | 1614 | 100.0% |

**Supplementary Table S4. Complete gene list. homology and GO**

| Tags | SeqName | Description | Length | #Hits | e-Value | sim mean | #GO | GO IDs | GO Names | Enzyme Codes | Enzyme Names | InterPro IDs | InterPro GO | InterPro GO Names |
|---|---|---|---|---|---|---|---|---|---|---|---|---|---|---|
| [INTERPRO, BLASTED] | Eg01_t000010 | PREDICTED: uncharacterized protein LOC108952056 | 147 | 6 | 1.75E-39 | 77.97 | | | | | | no IPS match | no IPS match | no IPS match |
| [INTERPRO, BLASTED, | Eg01_t000020 | conserved oligomeric Golgi complex subunit 6 | 101 | 10 | 3.44E-51 | 91.34 | 4 | P:GO:0006891 | P:intra-Golgi vesicle-mediated transpo | | | IPR010490 (PFAM | P:GO:0006891 | P:intra-Golgi vesicle-mediated transport; C:Golgi transport complex |
| [INTERPRO, BLASTED, | Eg01_t000030 | PREDICTED: uncharacterized protein LOC103988795 | 430 | 10 | 0 | 93.23 | 4 | P:GO:0001522 | P:pseudourid | EC:3.2; EC:4 | Glycosylases | IPR019339 (SMAR | F:GO:0016798 | F:hydrolase activity, acting on glycosyl bonds |
| [INTERPRO, BLASTED, | Eg01_t000040 | arabinosyltransferase XEG113 isoform X1 | 268 | 10 | 6.84E-119 | 86.98 | 1 | C:GO:0016021 | C:integral component of membrane | | | IPR005069 (PFAM | no GO terms | no GO terms |
| [INTERPRO, BLASTED, | Eg01_t000050 | arabinosyltransferase XEG113 isoform X2 | 327 | 10 | 1.05E-160 | 69.09 | 1 | C:GO:0016020 | C:membrane | | | IPR005069 (PFAM | no GO terms | no GO terms |
| [INTERPRO, NO-BLAST] | Eg01_t000060 | ---NA--- | 221 | | | | | | | | | no IPS match | no IPS match | no IPS match |
| [INTERPRO, BLASTED, | Eg01_t000070 | ubiquitin carboxyl-terminal hydrolase 2 | 334 | 10 | 0 | 94.86 | 3 | P:GO:0006511 | P:ubiquitin- | EC:3.4.19.12 | Ubiquitinyl | IPR001578 (PRIN | P:GO:0006511 | P:ubiquitin-dependent protein catabolic process; F:thiol-dependent deubiquitinase |
| [INTERPRO, BLASTED, | Eg01_t000080 | protein BTR1-like | 339 | 10 | 0 | 92.6 | 3 | P:GO:0010468 | P:regulation of gene expression; F:mRN | | | IPR004087 (SMAR | F:GO:0003676 | F:nucleic acid binding; F:RNA binding |
| [INTERPRO, BLASTED, | Eg01_t000090 | nuclear transcription factor Y subunit C-1-like | 106 | 10 | 2.83E-56 | 85.81 | 4 | P:GO:0045944 | P:positive regulation of transcription | | | IPR003958 (PFAM | no GO terms | no GO terms |
| [INTERPRO, NO-BLAST] | Eg01_t000100 | ---NA--- | 192 | | | | | | | | | no IPS match | no IPS match | no IPS match |
| [INTERPRO, NO-BLAST] | Eg01_t000110 | ---NA--- | 104 | | | | | | | | | no IPS match | no IPS match | no IPS match |
| [INTERPRO, BLASTED, | Eg01_t000120 | protein odr-4 homolog | 496 | 10 | 0 | 81.01 | 3 | P:GO:0055085 | P:transmembr | EC:7 | Translocases | IPR029454 (PFAM | no GO terms | no GO terms |
| [INTERPRO, BLASTED, | Eg01_t000130 | choline transporter-like protein 2 isoform X1 | 709 | 10 | 0 | 95.08 | 3 | P:GO:0055085 | P:transmembr | EC:7 | Translocases | IPR007603 (PFAM | no GO terms | no GO terms |
| [INTERPRO, BLASTED] | Eg01_t000140 | hypothetical protein BHE74_00035225 | 165 | 1 | 1.01E-27 | 57.93 | | | | | | no IPS match | no IPS match | no IPS match |
| [INTERPRO, BLASTED] | Eg01_t000150 | PREDICTED: uncharacterized protein LOC103977312 | 318 | 10 | 0 | 91.61 | | | | | | IPR021920 (PFAM | no GO terms | no GO terms |
| [INTERPRO, NO-BLAST] | Eg01_t000160 | ---NA--- | 139 | | | | | | | | | no IPS match | no IPS match | no IPS match |
| [INTERPRO, BLASTED] | Eg01_t000170 | UPF0051 protein in atpA 3'region-like | 57 | 5 | 9.08E-31 | 93.33 | | | | | | no IPS match | no IPS match | no IPS match |
| [INTERPRO, BLASTED, | Eg01_t000180 | PREDICTED: uncharacterized protein LOC108952358 | 407 | 10 | 0 | 78.95 | 4 | P:GO:0006355 | P:regulation of transcription, DNA-tem | | | IPR003340 (SMAR | F:GO:0003677 | F:DNA binding |
| [INTERPRO, BLASTED, | Eg01_t000190 | dihydroorotate dehydrogenase (quinone), mitochon | 460 | 10 | 0 | 93.55 | 4 | P:GO:0006207 | P:'de novo' | EC:1.3.5.2 | Dihydroorota | IPR005720 (PFAM | P:GO:0055114 | P:obsolete oxidation-reduction process; F:oxidoreductase activity, acting on the CH-CH group of donors; C:cytoplasm |
| [INTERPRO, BLASTED, | Eg01_t000200 | dolichyl-diphosphooligosaccharide--protein glyco | 730 | 10 | 0 | 95.12 | 4 | P:GO:0006486 | P:protein gl | EC:2.4.99.18 | Dolichyl-dip | IPR003674 (PFAM | P:GO:0006486 | P:protein glycosylation; F:oligosaccharyl transferase activity; C:membrane |
| [INTERPRO, BLASTED] | Eg01_t000210 | PREDICTED: uncharacterized protein LOC103977307 | 304 | 10 | 0 | 83.99 | | | | | | IPR022228 (PFAM | no GO terms | no GO terms |
| [INTERPRO, BLASTED, | Eg01_t000220 | TSL-kinase interacting protein 1 | 821 | 10 | 0 | 78.38 | 2 | P:GO:0016310 | P:phosphorylation; F:kinase activity | | | IPR001005 (SMAR | no GO terms | no GO terms |
| [INTERPRO, BLASTED] | Eg01_t000230 | pentatricopeptide repeat-containing protein At3g | 410 | 10 | 0 | 93.06 | | | | | | IPR002885 (PFAM | no GO terms | no GO terms |
| [INTERPRO, NO-BLAST] | Eg01_t000240 | ---NA--- | 92 | | | | | | | | | no IPS match | no IPS match | no IPS match |
| [INTERPRO, BLASTED] | Eg01_t000250 | pentatricopeptide repeat-containing protein At1g | 575 | 10 | 0 | 91.78 | | | | | | IPR002885 (PFAM | no GO terms | no GO terms |
| [INTERPRO, BLASTED, | Eg01_t000260 | ethylene-responsive transcription factor 14-like | 121 | 10 | 9.06E-82 | 90.7 | 5 | P:GO:0006355 | P:regulation of transcription, DNA-tem | | | IPR001471 (PRIN | P:GO:0006355 | P:regulation of transcription, DNA-templated; F:DNA-binding transcription factor activity |
| [INTERPRO, BLASTED, | Eg01_t000270 | Cell division cycle protein | 203 | 10 | 1.07E-99 | 73.34 | 2 | F:GO:0005524 | F:ATP binding; F:ATP hydrolysis activi | | | IPR003959 (PFAM | F:GO:0005524 | F:ATP binding; F:ATP hydrolysis activity |
| [INTERPRO, BLASTED, | Eg01_t000280 | Cell division cycle protein 48 | 202 | 10 | 1.69E-89 | 82.49 | 7 | P:GO:0006259 | P:DNA metabo | EC:3 | Hydrolases | IPR004201 (SMAR | F:GO:0005524 | F:ATP binding; F:ATP hydrolysis activity |
| [INTERPRO, BLASTED, | Eg01_t000290 | U-box domain-containing protein 12 | 641 | 10 | 0 | 85.77 | 4 | P:GO:0007166 | P:cell surfa | EC:2 | Transferases | IPR000225 (SMAR | P:GO:0016567 | P:protein ubiquitination; F:ubiquitin-protein transferase activity; F:protein binding |
| [INTERPRO, BLASTED, | Eg01_t000300 | probable pectate lyase 8 isoform X2 | 577 | 10 | 0 | 92.53 | 3 | P:GO:0045490 | P:pectin cat | EC:4.2.2.2 | Pectate lyas | IPR018082 (PRIN | no GO terms | no GO terms |
| [INTERPRO, BLASTED, | Eg01_t000310 | 3-ketoacyl-CoA thiolase 2, peroxisomal-like | 364 | 10 | 0 | 77.14 | 4 | P:GO:0006635 | P:fatty acid | EC:2.3.1.16 | Acetyl-CoA C | IPR020617 (PFAM | F:GO:0016747 | F:acyltransferase activity, transferring groups other than amino-acyl groups |
| [INTERPRO, BLASTED, | Eg01_t000320 | dehydration-responsive element-binding protein 2 | 195 | 10 | 1.06E-137 | 85.33 | 4 | P:GO:0045893 | P:positive regulation of transcription | | | IPR001471 (PRIN | P:GO:0006355 | P:regulation of transcription, DNA-templated; F:DNA-binding transcription factor activity |
| [INTERPRO, NO-BLAST] | Eg01_t000330 | ---NA--- | 72 | | | | | | | | | no IPS match | no IPS match | no IPS match |
| [INTERPRO, BLASTED, | Eg01_t000340 | protein PAF1 homolog | 732 | 10 | 0 | 86 | 3 | P:GO:0006368 | P:transcription elongation from RNA po | | | IPR007133 (PFAM | P:GO:0006368 | P:transcription elongation from RNA polymerase II promoter; P:histone modification; C:Cdc73/Paf1 complex |
| [INTERPRO, BLASTED, | Eg01_t000350 | NAC domain-containing protein 14 isoform X1 | 723 | 10 | 0 | 76.86 | 3 | P:GO:0006355 | P:regulation of transcription, DNA-tem | | | IPR003441 (PFAM | P:GO:0006355 | P:regulation of transcription, DNA-templated; F:DNA binding |
| [INTERPRO, BLASTED, | Eg01_t000360 | hypothetical protein B296_00029256, partial | 155 | 1 | 1.88E-49 | 87 | 2 | P:GO:0006355 | P:regulation of transcription, DNA-tem | | | no IPS match | no IPS match | no IPS match |
| [INTERPRO, BLASTED, | Eg01_t000370 | TATA box-binding protein-associated factor RNA p | 792 | 10 | 0 | 77.76 | 4 | P:GO:0001188 | P:RNA polymerase I preinitiation compl | | | IPR021752 (PFAM | P:GO:0001188 | P:RNA polymerase I preinitiation complex assembly; P:transcription by RNA polymerase I; F:RNA polymerase I core promoter sequence-specific DNA binding; C:RNA polymerase I core factor complex |
| [INTERPRO, BLASTED, | Eg01_t000380 | transcription factor MYC2-like | 650 | 10 | 0 | 77.97 | 3 | P:GO:0006355 | P:regulation of transcription, DNA-tem | | | IPR011598 (SMAR | F:GO:0046983 | F:protein dimerization activity |
| [INTERPRO, NO-BLAST] | Eg01_t000390 | ---NA--- | 131 | | | | | | | | | no IPS match | no IPS match | no IPS match |
| [INTERPRO, BLASTED, | Eg01_t000400 | probable transcription factor KAN2 | 313 | 10 | 0 | 75.23 | 4 | P:GO:0006355 | P:regulation of transcription, DNA-tem | | | IPR001005 (PFAM | no GO terms | no GO terms |

TRUNCATED IN PDF VERSION

36796 ADDITIONAL LINES DELETED

**Supplementary Table S5. Statistics for shared orthogroups (OG) and gene clustering among E. glaucum, E. ventricosum, Musa acuminata, M. balbisiana, and M. schizocarpa genomes.**

| | *E. glaucum* | *E. ventricosum* | *M. acuminata* | *M. balbisiana* | *M. schizocarpa* |
|---|---|---|---|---|---|
| Number of genes | 36836 | 58438 | 35276 | 33021 | 32809 |
| Number of genes in orthogroups | 31450 | 47487 | 34006 | 29051 | 31031 |
| Number of unassigned genes | 5386 | 10951 | 1270 | 3970 | 1778 |
| Percentage of genes in orthogroups | 85.4 | 81.3 | 96.4 | 88 | 94.6 |
| Percentage of unassigned genes | 14.6 | 18.7 | 3.6 | 12 | 5.4 |
| Number of orthogroups containing species | 22490 | 23583 | 23928 | 20441 | 22635 |
| Percentage of orthogroups containing species | 74.6 | 78.3 | 79.4 | 67.8 | 75.1 |
| Number of species-specific orthogroups | 115 | 1466 | 41 | 170 | 53 |
| Number of genes in species-specific orthogroups | 276 | 8365 | 109 | 771 | 151 |
| Percentage of genes in species-specific orthogroups | 0.7 | 14.3 | 0.3 | 2.3 | 0.5 |

1) E. ventricosum annotation on draft genome assembly over-representing number of genes (58438); other species based on chromosome-level assemblies

**Supplementary table S6. Positively selected genes and their annotation**

| EGL gene | MAC gene | Ka | Ks | Ka/Ks | P.Value. | Tags | Description | Length | #Hits | e-Value | sim mear | #GO | GO IDs | GO Name: | Enzyme ( | Enzyme N | InterPro IDs | InterPro | InterPro GO Names |
|---|---|---|---|---|---|---|---|---|---|---|---|---|---|---|---|---|---|---|---|
| Eg01_t016530 | Macma4_04_g20080 | 0.03793 | 7.59E-04 | 50 | 0.12055 | [INTERPI | putative FCS-Like Zinc finger 3 | 120 | 10 | 7.55E-72 | 78.18 | NA | NA | NA | NA | NA | IPR007650 (PFAM); PTHR | no GO te | no GO terms |
| Eg07_t043580 | Macma4_10_g02680 | 0.03071 | 0.000614 | 50 | 0.20224 | [INTERPI | non-specific lipid-transfer protein 1-like | 112 | 5 | 2.63E-23 | 57.41 | 2 | P:GO:00( | P:lipid | NA | NA | IPR000528 (PRINTS); IF | P:GO:00( | P:lipid transport; F:lipid binding |
| Eg03_t017210 | Macma4_03_g32730 | 0.02366 | 4.74E-04 | 49.9537 | 0.4666 | [INTERPI | ---NA--- | 85 | NA | NA | NA | NA | NA | NA | NA | NA | no IPS match | no IPS n | no IPS match |
| Eg05_t023490 | Macma4_05_g22650 | 0.13947 | 0.032455 | 4.29722 | 0.00366 | [INTERPI | transcription factor DIVARICATA-like isoform X1 | 118 | 10 | 1.56E-59 | 71.3 | 2 | P:GO:00( | P:regula | NA | NA | IPR001005 (SMART); IPF | no GO te | no GO terms |
| Eg04_t033180 | Macma4_04_g05850 | 0.0589 | 0.014442 | 4.0786 | 0.18456 | [INTERPI | hypothetical protein C4D60_Mb04t05480 | 99 | 10 | 2.85E-48 | 81.51 | 2 | P:GO:004 | P:phospl | NA | NA | PTHR35282 (PANTHER); F | no GO te | no GO terms |
| Eg07_t000020 | Macma4_09_g23170 | 0.05374 | 0.015007 | 3.58107 | 0.12922 | [INTERPI | hypothetical protein GW17_00041715 | 95 | 10 | 6.09E-58 | 77.17 | NA | NA | NA | NA | NA | PTHR35162 (PANTHER); F | no GO te | no GO terms |
| Eg09_t003020 | Macma4_11_g07000 | 0.04203 | 0.011812 | 3.55804 | 0.1094 | [INTERPI | ---NA--- | 103 | NA | NA | NA | NA | NA | NA | NA | NA | IPR039618 (PANTHER) | no GO te | no GO terms |
| Eg09_t021450 | Macma4_11_g04160 | 0.08995 | 0.026656 | 3.37434 | 0.03312 | [INTERPI | autophagy-related protein 8f | 130 | 5 | 2.35E-57 | 81.95 | 2 | P:GO:00( | P:autopl | NA | NA | IPR004241 (PFAM); IPR( | no GO te | no GO terms |
| Eg06_t002960 | Macma4_09_g19460 | 0.06156 | 0.018545 | 3.31973 | 0.09269 | [INTERPI | uncharacterized protein LOC109506234 | 71 | 5 | 3.86E-26 | 84.33 | 1 | C:GO:001 | C:integi | NA | NA | PTHR37225 (PANTHER) | no GO te | no GO terms |
| Eg06_t014530 | Macma4_07_g09960 | 0.06496 | 0.019736 | 3.29138 | 0.08812 | [INTERPI | uncharacterized protein At2g34160 | 128 | 5 | 8.04E-67 | 88.21 | 2 | F:GO:00( | F:RNA b: | NA | NA | IPR002775 (PFAM); IPR( | F:GO:00( | F:nucleic acid binding |
| Eg07_t003180 | Macma4_07_g20430 | 0.04977 | 0.015284 | 3.25628 | 0.25606 | [INTERPI | hypothetical protein C4D60_Mb07t09140 | 85 | 10 | 1.19E-44 | 75.95 | 1 | C:GO:001 | C:membra | NA | NA | PTHR37908 (PANTHER) | no GO te | no GO terms |
| Eg05_t015980 | Macma4_05_g15890 | 0.06717 | 0.020962 | 3.20441 | 0.03673 | [INTERPI | ras-related protein RABC2a | 209 | 5 | 2.99E-98 | 80.37 | 4 | P:GO:00( | P:intrac | EC:3.6.1 | Acting ( | PR00449 (PRINTS); SMO( | F:GO:00( | F:GTPase activity; F:GTP binding |
| Eg09_t022790 | Macma4_11_g03020 | 0.02575 | 0.008123 | 3.16947 | 0.32147 | [INTERPI | uncharacterized protein LOC117906292 | 118 | 5 | 5.64E-53 | 83.92 | 1 | C:GO:001 | C:integi | NA | NA | IPR000620 (PFAM); IPR( | C:GO:001 | C:membrane; C:integral component of membrane |
| Eg06_t001300 | Macma4_Mt_g00610 | 0.03784 | 0.013267 | 2.8521 | 0.18621 | [INTERPI | cytochrome c biogenesis FN (mitochondrion) | 108 | 5 | 5.25E-56 | 89.23 | 6 | P:GO:001 | P:heme t | EC:7 | Transloc | IPR003569 (PRINTS); PT | P:GO:001 | P:heme transport; P:cytochrome complex assembly; F:heme transmembrane transporter activity; C:membrane |
| Eg01_t006570 | Macma4_03_g05980 | 0.04967 | 0.01805 | 2.75159 | 0.40678 | [INTERPI | dormancy-associated protein homolog 4-like | 97 | 10 | 2.50E-48 | 76.59 | NA | NA | NA | NA | NA | IPR008406 (PANTHER); F | no GO te | no GO terms |
| Eg04_t008800 | Macma4_06_g37820 | 0.12868 | 0.047102 | 2.73185 | 0.06977 | [INTERPI | PREDICTED: uncharacterized protein LOC10399026 | 98 | 1 | 3.34E-25 | 81.82 | 1 | P:GO:003 | P:regula | NA | NA | IPR040389 (PANTHER); F | P:GO:003 | P:regulation of DNA endoreduplication |
| Eg04_t031890 | Macma4_04_g07080 | 0.03513 | 0.013163 | 2.66883 | 0.36681 | [INTERPI | photosystem II 5 kDa protein, chloroplastic-lik | 104 | 10 | 2.34E-70 | 81.29 | NA | NA | NA | NA | NA | IPR040296 (PANTHER) | no GO te | no GO terms |
| Eg05_t029510 | Macma4_05_g32810 | 0.12718 | 0.048371 | 2.6292 | 0.01555 | [INTERPI | ---NA--- | 112 | NA | NA | NA | NA | NA | NA | NA | NA | no IPS match | no IPS n | no IPS match |
| Eg03_t009880 | Macma4_03_g25710 | 0.10739 | 0.042993 | 2.49793 | 0.00163 | [INTERPI | uncharacterized protein LOC105036081 | 313 | 5 | 4.24E-77 | 52.82 | 2 | C:GO:001 | C:membra | NA | NA | PTHR33625:SF4 (PANTHEF | no GO te | no GO terms |
| Eg07_t029320 | Macma4_09_g13060 | 0.11084 | 0.045303 | 2.44671 | 0.04286 | [INTERPI | GPI-anchored protein LLG1-like | 168 | 5 | 2.21E-56 | 74.41 | 2 | C:GO:001 | C:membra | NA | NA | IPR039307 (PANTHER); F | no GO te | no GO terms |
| Eg01_t026170 | Macma4_01_g19590 | 0.08955 | 0.0374 | 2.39427 | 0.14861 | [INTERPI | hypothetical protein B296_00059089, partial | 113 | 9 | 3.89E-68 | 80.98 | 1 | C:GO:001 | C:membra | NA | NA | no IPS match | no IPS n | no IPS match |
| Eg09_t024580 | Macma4_11_g01300 | 0.09004 | 0.0386 | 2.33258 | 0.16074 | [INTERPI | succinate dehydrogenase subunit 3-2, mitochondı | 123 | 5 | 5.71E-23 | 67.68 | 3 | P:GO:00( | P:aerob: | NA | NA | PTHR10978 (PANTHER); F | no GO te | no GO terms |
| Eg07_t024440 | Macma4_09_g20200 | 0.05941 | 0.026829 | 2.21423 | 0.35469 | [INTERPI | putative Triosephosphate isomerase | 106 | 10 | 2.24E-52 | 78.42 | NA | NA | NA | NA | NA | PTHR36385 (PANTHER) | no GO te | no GO terms |
| Eg06_t022460 | Macma4_07_g02410 | 0.17301 | 0.079827 | 2.16731 | 0.00396 | [INTERPI | uncharacterized protein LOC105032758 | 225 | 5 | 4.04E-61 | 66.26 | 2 | C:GO:001 | C:membra | NA | NA | PTHR35508 (PANTHER); F | no GO te | no GO terms |
| Eg04_t001070 | Macma4_04_g04020 | 0.13695 | 0.063679 | 2.15068 | 0.14555 | [INTERPI | hypothetical protein C4D60_Mb04t03730 | 76 | 7 | 1.32E-34 | 79.7 | 3 | P:GO:005 | P:transı | EC:7 | Transloc | no IPS match | no IPS n | no IPS match |
| Eg01_t017860 | Macma4_04_g21230 | 0.09768 | 0.045729 | 2.13597 | 0.00318 | [INTERPI | hypothetical protein GW17_00056264, partial | 372 | 2 | 2.21E-118 | 93.87 | NA | NA | NA | NA | NA | PR01228 (PRINTS) | no GO te | no GO terms |
| Eg03_t014420 | Macma4_03_g29940 | 0.13488 | 0.063482 | 2.12462 | 0.00119 | [INTERPI | ---NA--- | 361 | NA | NA | NA | NA | NA | NA | NA | NA | no IPS match | no IPS n | no IPS match |
| Eg02_t034660 | Macma4_02_g14110 | 0.11545 | 0.054474 | 2.11935 | 0.05567 | [INTERPI | E3 ubiquitin-protein ligase RING1-like | 196 | 10 | 1.36E-126 | 71.52 | 4 | P:GO:001 | P:monosa | EC:7 | Transloc | IPR001841 (SMART); IPF | no GO te | no GO terms |
| Eg03_t031130 | Macma4_08_g24200 | 0.22184 | 0.105611 | 2.10051 | 0.01668 | [INTERPI | heavy metal-associated isoprenylated plant prot | 195 | 5 | 2.07E-22 | 69.66 | 1 | F:GO:004 | F:metal | NA | NA | PTHR45811 (PANTHER); F | no GO te | no GO terms |
| Eg01_t027720 | Macma4_01_g11880 | 0.10224 | 0.048718 | 2.09867 | 0.031 | [INTERPI | zinc finger protein ZAT2-like | 258 | 10 | 8.05E-114 | 81.53 | 2 | P:GO:00( | P:regula | NA | NA | IPR013087 (SMART); PF1 | no GO te | no GO terms |
| Eg08_t016400 | Macma4_10_g19800 | 0.04871 | 0.023973 | 2.03199 | 0.09933 | [INTERPI | transcription factor VIP1 | 325 | 5 | 6.52E-119 | 67.11 | 2 | P:GO:00( | P:regula | NA | NA | IPR004827 (SMART); IPF | P:GO:00( | P:regulation of transcription, DNA-templated; F:DNA-binding transcription factor activity |
| Eg06_t011770 | Macma4_07_g14070 | 0.03945 | 0.019536 | 2.01946 | 0.16642 | [INTERPI | nudix hydrolase 15, mitochondrial-like | 269 | 5 | 2.97E-122 | 74.52 | 1 | F:GO:001 | F:CoA py | EC:3.6.1 | Acting ( | IPR000086 (PFAM); PTHF | F:GO:001 | F:hydrolase activity |
| Eg02_t041650 | Macma4_02_g25540 | 0.03078 | 0.015539 | 1.98048 | 0.55743 | [INTERPI | hypothetical protein BHM03_00045241 | 80 | 5 | 1.87E-48 | 75.89 | NA | NA | NA | NA | NA | PTHR37908 (PANTHER) | no GO te | no GO terms |
| Eg01_t020940 | Macma4_06_g40300 | 0.11951 | 0.06093 | 1.96145 | 0.47978 | [INTERPI | hypothetical chloroplast RF19 | 78 | 10 | 1.67E-45 | 92.15 | 3 | P:GO:001 | P:prote: | NA | NA | no IPS match | no IPS n | no IPS match |
| Eg01_t034980 | Macma4_01_g10820 | 0.05673 | 0.029882 | 1.89858 | 0.51131 | [INTERPI | PREDICTED: uncharacterized protein LOC10398362 | 98 | 10 | 1.71E-57 | 71.77 | 1 | C:GO:001 | C:membra | NA | NA | PTHR37741 (PANTHER) | no GO te | no GO terms |
| Eg08_t046190 | Macma4_06_g12990 | 0.06477 | 0.034217 | 1.89288 | 0.11697 | [INTERPI | zinc finger protein 4 | 221 | 5 | 4.09E-74 | 71.3 | NA | NA | NA | NA | NA | PTHR45730:SF44 (PANTHE | no GO te | no GO terms |
| Eg07_t019370 | Macma4_09_g29390 | 0.04752 | 0.025181 | 1.88718 | 0.59037 | [INTERPI | heavy metal-associated isoprenylated plant prot | 89 | 10 | 2.91E-51 | 88.83 | 1 | F:GO:004 | F:metal | NA | NA | PTHR47294 (PANTHER); F | no GO te | no GO terms |
| Eg06_t020970 | Macma4_07_g03910 | 0.0671 | 0.035802 | 1.87418 | 0.2813 | [INTERPI | GATA transcription factor 16 | 130 | 5 | 3.40E-28 | 62.25 | 3 | P:GO:00( | P:regula | NA | NA | IPR000679 (SMART); IPF | P:GO:00( | P:regulation of transcription, DNA-templated; F:zinc ion binding; F:sequence-specific DNA binding |
| Eg07_t014790 | Macma4_09_g24810 | 0.03004 | 0.016192 | 1.85511 | 0.55837 | [INTERPI | hypothetical protein BHE74_00023195 | 91 | 10 | 1.72E-61 | 80.13 | NA | NA | NA | NA | NA | PTHR33181:SF19 (PANTHE | no GO te | no GO terms |
| Eg04_t005980 | Macma4_06_g40110 | 0.0569 | 0.030744 | 1.85087 | 0.4067 | [INTERPI | cytochrome c oxidase assembly protein cox16, mi | 132 | 10 | 6.66E-86 | 90.03 | 2 | C:GO:00( | C:mitocl | NA | NA | IPR020164 (PFAM) | C:GO:003 | C:mitochondrial membrane |

287 ADDITIONAL LINES DELETED FROM BIORXIV TABLE

Truncated in PDF Version ···
288 ADDITIONAL LINES DELETED

**Supplementary Table S7: Result of gene family size change analysis.**

| Species | Expanded families | Genes gained | genes/ expansion | Contracted families | Genes lost | genes/ contraction | Unchanged | Avgerage expansion |
|---|---|---|---|---|---|---|---|---|
| *M. balbisiana* | 550 (42) | 1246 | 2.27 | 2670 (59) | 3114 | 1.17 | 9164 | -0.15084 |
| *M. schizocarpa* | 573 (21) | 654 | 1.14 | 1139 (74) | 1397 | 1.23 | 10672 | -0.0599968 |
| *E. glaucum* | 1229 (25) | 1533 | 1.25 | 837 (29) | 904 | 1.08 | 10318 | 0.0507913 |
| *M. acuminata* | 872 (70) | 1247 | 1.43 | 400 (23) | 442 | 1.1 | 11112 | 0.0650032 |
| *Phoenix dactylifera* | 2009 (29) | 2984 | 1.49 | 3444 (35) | 4271 | 1.24 | 6931 | -0.103924 |
| *Oryza sativa* | 1498 (79) | 2907 | 1.94 | 2793 (3) | 3512 | 1.26 | 8093 | -0.0488534 |

### Supplementary Table S8 - a) molecular function (MF); b) biological process (BP). (See Fig. 3B)

#### a) Top 20 GO molecular function enrichments for E. glaucum and shared E. glaucum/E. ventricosum gene families

| GO ID | Term | Annotated | Significant | Expected | classicFisher |
|---|---|---|---|---|---|
| GO:0005506 | iron ion binding | 295 | 13 | 3.48 | 5.60E-09 |
| GO:0005515 | protein binding | 3181 | 37 | 37.56 | 2.80E-05 |
| GO:0004190 | aspartic-type endopeptidase activity | 89 | 5 | 1.05 | 0.0001 |
| GO:0003849 | 3-deoxy-7-phosphoheptulonate synthase | 6 | 2 | 0.07 | 0.00039 |
| GO:0003677 | DNA binding | 1130 | 19 | 13.34 | 0.0005 |
| GO:0022857 | transmembrane transporter activity | 871 | 21 | 10.28 | 0.00064 |
| GO:0016787 | hydrolase activity | 1977 | 29 | 23.34 | 0.00097 |
| GO:0015165 | pyrimidine nucleotide-sugar transmembrane | 11 | 2 | 0.13 | 0.00141 |
| GO:0004672 | protein kinase activity | 959 | 13 | 11.32 | 0.00154 |
| GO:0016747 | transferase activity, transferring acyl | 245 | 6 | 2.89 | 0.00187 |
| GO:0015211 | purine nucleoside transmembrane transporter | 14 | 2 | 0.17 | 0.00231 |
| GO:0008686 | 3,4-dihydroxy-2-butanone-4- phosphate syn | 1 | 1 | 0.01 | 0.00515 |
| GO:0030170 | pyridoxal phosphate binding | 21 | 2 | 0.25 | 0.0052 |
| GO:0003975 | UDP-N-acetylglucosamine-dolichyl-phospha | 2 | 1 | 0.02 | 0.01028 |
| GO:0003979 | UDP-glucose 6-dehydrogenase activity | 2 | 1 | 0.02 | 0.01028 |
| GO:0052861 | glucan endo-1,3-beta-glucanase activity | 2 | 1 | 0.02 | 0.01028 |
| GO:0004540 | ribonuclease activity | 32 | 2 | 0.38 | 0.01185 |
| GO:0015930 | glutamate synthase activity | 3 | 1 | 0.04 | 0.01538 |
| GO:0060090 | molecular adaptor activity | 3 | 1 | 0.04 | 0.01538 |
| GO:0030599 | pectinesterase activity | 38 | 2 | 0.45 | 0.01645 |

#### b) Top 20 GO biological process enrichments for E. glaucum and shared E. glaucum/E. ventricosum gene families

| GO ID | Term | Annotated | Significant | Expected | classicFisher |
|---|---|---|---|---|---|
| GO:0006355 | regulation of transcription | 178 | 14 | 2.37 | 8.80E-10 |
| GO:0032875 | regulation of DNA endoreduplication | 13 | 3 | 0.17 | 0.00024 |
| GO:0000160 | phosphorelay signal transduction system | 53 | 4 | 0.7 | 0.00173 |
| GO:0055085 | transmembrane transport | 159 | 7 | 2.11 | 0.00217 |
| GO:0009765 | photosynthesis, light harvesting | 29 | 3 | 0.39 | 0.00277 |
| GO:0015074 | DNA integration | 1 | 1 | 0.01 | 0.00987 |
| GO:0098869 | cellular oxidant detoxification | 3 | 1 | 0.04 | 0.02934 |
| GO:0006120 | mitochondrial electron transport, NADH t... | 4 | 1 | 0.05 | 0.03893 |
| GO:0016560 | protein import into peroxisome matrix, d... | 4 | 1 | 0.05 | 0.03893 |
| GO:0006952 | defense response | 61 | 3 | 0.81 | 0.04506 |
| GO:0051225 | spindle assembly | 7 | 1 | 0.09 | 0.06715 |
| GO:0009966 | regulation of signal transduction | 8 | 1 | 0.11 | 0.07637 |
| GO:0051513 | regulation of monopolar cell growth | 12 | 1 | 0.16 | 0.11238 |
| GO:0046622 | positive regulation of organ growth | 13 | 1 | 0.17 | 0.12116 |
| GO:0008299 | isoprenoid biosynthetic process | 13 | 1 | 0.17 | 0.12116 |
| GO:0010112 | regulation of systemic acquired resistance | 14 | 1 | 0.19 | 0.12986 |
| GO:0009734 | auxin-activated signaling pathway | 14 | 1 | 0.19 | 0.12986 |
| GO:0009639 | response to red or far red light | 17 | 1 | 0.23 | 0.15546 |
| GO:0006281 | DNA repair | 40 | 1 | 0.53 | 0.32863 |
| GO:0000398 | mRNA splicing, via spliceosome | 44 | 1 | 0.58 | 0.35496 |

**Supplementary Table S9. Comparison of transcription factors between Ensete glaucum and Musa acuminata**

| TF family | *Ensete glaucum* | *Musa acuminata* |
|---|---|---|
| AP2/ERF-AP2 | 37 | 72 |
| AP2/ERF-ERF | 237 | 264 |
| AP2/ERF-RAV | 11 | 23 |
| B3 | 46 | 58 |
| B3-ARF | 60 | 111 |
| BBR-BPC | 11 | 32 |
| BES1 | 10 | 13 |
| bHLH | 248 | 445 |
| bZIP | 138 | 232 |
| CO-like | 14 | 24 |
| C2C2-Dof | 85 | 111 |
| GATA | 48 | 74 |
| C2C2-LSD | 8 | 13 |
| YABBY | 22 | 25 |
| C2H2 | 191 | 237 |
| C3H | 64 | 116 |
| CAMTA | 4 | 31 |
| CPP | 9 | 12 |
| DBB | 8 | 24 |
| E2F-DP | 11 | 21 |
| EIL | 17 | 20 |
| FAR1 | 44 | 96 |
| GARP-ARR-B | 13 | 23 |
| G2-like | 101 | 163 |
| GeBP | 11 | 14 |
| GRAS | 68 | 94 |
| GRF | 16 | 30 |
| HB-HD-ZIP | 69 | 117 |
| HB-KNOX | 15 | |
| HB-other | 27 | 19 |
| HB-PHD | 2 | 5 |
| HB-WOX | 26 | 30 |
| HRT | 1 | 1 |
| HSF | 43 | 55 |
| LFY | 1 | 1 |
| MADS-MIKC | 28 | 92 |
| MADS-M-type | 41 | 32 |
| MYB | 260 | 363 |
| MYB-related | 109 | 141 |
| NAC | 202 | 187 |
| NF-X1 | 1 | 2 |
| NF-YA | 19 | 39 |
| NF-YB | 21 | 24 |
| NF-YC | 16 | 39 |
| S1Fa-like | 2 | 4 |
| SAP | 3 | 4 |
| SBP | 63 | 97 |
| SRS | 8 | 12 |
| STAT | 2 | 1 |
| TCP | 45 | 63 |
| Trihelix | 55 | 75 |
| VOZ | 3 | 5 |
| Whirly | 3 | 4 |
| WRKY | 159 | 215 |
| zf-HD | 27 | 36 |

**Supplementary Table S10. Transposable elements and other repeat proportions comparison in assembly (RepeatMasker).**

| | | | *Ensete glaucum* | *Musa acuminata* |
|---|---|---|---|---|
| LTR retroelements | Ty1/Copia | Ale | 2.26% | 0.73% |
| | | Alesia | 0.03% | 0.05% |
| | | Angela | 4.67% | 4.12% |
| | | Ikeros | 1.00% | 1.13% |
| | | Ivana | 1.17% | 0.46% |
| | | SIRE | 5.34% | 18.68% |
| | | TAR | 0.09% | 0.10% |
| | | Tork | 1.28% | 1.55% |
| | | Unclassified | 0.90% | 2.03% |
| | | Total Copia | 17.64% | 28.86% |
| | Ty3/Gypsy | CRM | 0.60% | 0.74% |
| | | Galadriel | 1.89% | 0.96% |
| | | Reina | 10.88% | 4.20% |
| | | Retand | 5.44% | 1.58% |
| | | Tekay | 0.34% | 1.62% |
| | | Unclassified Gypsy | 0.11% | 1.89% |
| | | Total Gypsy | 19.25% | 10.98% |
| | Unclassified LTR | | 0.31% | 0.20% |
| Other Class I | LINE | | 0.77% | 2.03% |
| DNA transposon | | hAT | 1.32% | 0.59% |
| | | CACTA | 1.24% | 0.31% |
| | | PIF-Harbinger | 0.39% | 0.14% |
| | | Mutator | 2.73% | 0.59% |
| | | Tc1-Mariner | 0.31% | 0.06% |
| | | Helitron | 1.19% | 0.25% |
| | | Total DNA transposon | 7.18% | 1.93% |
| Unclassfied | | | 8.74% | 9.56% |
| | | Interspersed repeats | 53.89% | 53.55% |
| | | Low_complexity | 0.16% | 0.14% |
| | | Simple_repeat | 0.97% | 1.29% |

**Supplementary Table S11. Repeat content (RepeatExplorer, %) comparison between different Musaceae genomes**

| Repeat | | Lineage/class | Clade | alternative names | *Musa acuminata* | *Musa balbisiana* | *Musa schizocarpa* | *Ensete ventricosum* | *Ensete glaucum* | *Musella lasiocarpa* |
|---|---|---|---|---|---|---|---|---|---|---|
| LTR retroelements | Ty1/Copia | Maximus-SIRE | | | # | 5.962 | 7.326 | 2.769 | # | 3.15 |
| | | Angela | | | # | 2.812 | 4.5 | 3.423 | 3.76 | # |
| | | Tork | | Tnt | # | 1.432 | 0.305 | 1.46 | # | # |
| | | Ale | | Hopscotch | # | 0.579 | 0.293 | 0.923 | # | # |
| | | Ivana | | Oryco | # | 0.181 | 1.491 | 0.967 | # | # |
| | | TAR | | Tont | # | 0.043 | | | # | |
| | | Ikeros | | | # | 0.424 | 0.308 | 0.505 | # | # |
| | | Bianca | | | | | 0.271 | | | |
| | | Total Ty1/Copia | | | # | # | # | 10.047 | | # |
| | Ty3/Gypsy | Chromovirus | Reina | | # | 3.134 | 0.372 | 5.386 | # | # |
| | | | Tekay | | # | 0.977 | 1.713 | 0.327 | # | # |
| | | | Galadriel | Monkey | # | 0.716 | | 1.823 | # | # |
| | | | CRM | | # | 0.379 | 0.023 | 0.56 | # | # |
| | | | Retand | | | 0.014 | 0.841 | 0.678 | # | 0.01 |
| | | | unclassified | | | | | | | |
| | | Total Ty3/Gypsy | | | # | 5.22 | 2.949 | 8.774 | # | # |
| | Unclassified LTR | | | | # | | 4.105 | | | |
| Other | Other Class_I pararetrovirus | | | | # | 0.027 | | | | |
| | LINE | | | | 0.53 | 0.537 | | 0.689 | # | # |
| | DNA transposon | | | | 0.07 | 0.056 | 0.242 | 0.063 | # | # |
| | Tandem Repeats | rDNA | | | # | 1.477 | 2.426 | 0.602 | # | # |
| | | Satellites | | | # | 0.853 | 1.545 | 1.323 | # | # |
| Annotated repetitive total | | | | | # | # | # | 21.498 | # | # |
| Unclassified repetitive | | | | | # | # | # | 13.411 | # | # |
| All repetitive total | | | | | # | # | # | 34.909 | # | # |
| Percentage of reads in top clusters | | | | | 43 | 46 | 44 | 33 | 38 | 35 |
| unknown low copy | | | | | # | # | # | 65.091 | # | # |

Supplementary Table S12. Abundance of major tandemly repeated DNA repeats in Illumina raw reads

| | | | | | |
|---|---|---|---|---|---|
| Illumina sequencing | 245,852,534 | | 150 bp paired | Total base pairs | 36,877,880,100 |
| Genome size (contig length) | 495,175,598 | bp | | Coverage | 74 |
| 5S rDNA | 1,056 | bp monomer | | | |
| (5S rRNA-ITS) | 191,703 | reads map to monomer | | | |
| | 28,755,450 | bp of 5S rDNA sequenced | | | |
| | 0.078 | % of genome is 5S rDNA | | | |
| | 366 | copies of monomer | | | |
| 45S rDNA | 9,984 | bp monomer | | | |
| (18S-ITS1-5.8S-ITS2-26S-NTS) | 2,982,590 | reads map to monomer | | | |
| | 447,388,500 | bp of 45S rDNA sequenced | | | |
| | 1.213 | % of genome is 45S rDNA | | | |
| | 587 | copies of monomer | | | |
| Egcen | 134 | bp monomer | | | |
| | 1,764,271 | reads map to monomer | | | |
| | 264,640,650 | bp of Egcen sequenced | | | |
| | 0.718 | % of genome is Egcen | | | |
| | 26,518 | copies of monomer | | | |

**Supplementary Table S13. Inferred centromere mid positions from locations of interrupted tandem arrays of the EGcen centromeric sequence on the chromosome assemblies.**

| | inferred centomere mid position (nt) 1) | EGcen arrays | | |
|---|---|---|---|---|
| | | | start (nt) | end (nt) |
| **eg01** | **31,021,187** | left | 29,094,495 | 29,442,365 |
| | | right | 32,202,769 | 33,321,780 |
| **eg02** | | minor | 15,294,331 | 15,305,956 |
| | **20,260,322** | major | 20,159,964 | 20,380,714 |
| **eg03** | **22,080,873** | left | 20,270,153 | 20,673,355 |
| | | right | 23,487,807 | 24,012,165 |
| **eg04** | **7,241,115** | major | 6,801,269 | 7,545,391 |
| **eg05** | | minor | 16,472,716 | 16,505,318 |
| | | minor | 21,904,330 | 21,909,085 |
| | **29,209,736** | major | 28,888,983 | 29,576,579 |
| **eg06** | | minor | 9,790,449 | 10,183,543 |
| | | minor | 13,173,685 | 13,304,647 |
| | | minor | 15,220,287 | 15,431,266 |
| | **23,886,618** | major | 23,623,636 | 24,179,815 |
| **eg07** | **27,560,472** | left | 26,378,957 | 27,030,005 |
| | | right | 28,192,438 | 28,640,489 |
| **eg08** | **26,539,379** | major | 26,311,459 | 26,767,298 |
| **eg09** | **32,317,789** | left | 29,253,112 | 30,175,361 |
| | | right | 34,886,786 | 35,240,275 |

1) Midpoint between left and right arrays, or of the single major arrays

**Supplementary Table S14. Comparative survey of microsatellite sequences in Ensete glaucum genome with other sister species.**

| Species | *Ensete glaucum* | *Ensete ventricosum* | *Musa acuminata* | *Musa balbisiana* | *Musa itinerans* | *Musa schizocarpa* |
|---|---|---|---|---|---|---|
| Total number of sequences examined | 9 | 52522 | 24 | 12 | 75687 | 194 |
| Total size of examined sequences (bp) | 481507213 | 440718910 | 450848473 | 402547155 | 462139488 | 525283493 |
| Total number of identified microsatellites | 123884 | 102795 | 91882 | 83354 | 91589 | 114207 |
| Number of microsatellite containing sequences | 9 | 27000 | 24 | 12 | 7083 | 185 |
| Sequences contain more than 1 microsatellites | 9 | 17692 | 22 | 12 | 3803 | 161 |
| Microsatellites in compound formation | 3996 | 2361 | 3132 | 2840 | 2472 | 4546 |
| Microsatellites density (1 Microsatellites per ** bp) | 3886.7 | 4313.7 | 4906.8 | 4829.4 | 5045.8 | 4599.4 |
| Class I microsatellites | 56365 | 42368 | 44719 | 38950 | 41468 | 58596 |
| Class II microsatellites | 63523 | 58064 | 44031 | 41564 | 47649 | 51065 |
| Ratio Class I:ClassII | 0.9 | 0.7 | 1.0 | 0.9 | 0.9 | 1.1 |
| AT rich microsatellites | 83236 | 69000 | 57578 | 51071 | 58742 | 78651 |
| GC rich microsatellites | 15536 | 12720 | 12434 | 11467 | 10800 | 13027 |
| AT/GC balance microsatellites | 21116 | 18713 | 18738 | 17976 | 19575 | 17983 |
| Mono-nucleotide repeats | 37223 | 36397 | 20708 | 21342 | 19934 | 26052 |
| Di-nucleotide repeats | 48062 | 36002 | 45913 | 40503 | 43988 | 53456 |
| Tri-nucleotide repeats | 29547 | 24207 | 20064 | 17520 | 22130 | 26852 |
| Tera-nucleotide repeats | 3230 | 2363 | 2366 | 1846 | 2565 | 3562 |
| Penta-nucleotide repeats | 4458 | 2774 | 1579 | 1141 | 1562 | 1860 |
| Hexa-nucleotide repeats | 1364 | 1051 | 1252 | 1002 | 1410 | 2425 |

Data for Ensete ventricosum, Musa acuminata, Musa balbisiana, Musa itinerans and Musa schizocarpa from Biswas et al (2020)

| | |
|---|---|
| Total SSR (bp ) | 2850278 |
| Total genome size | 481507213 |
| % of SSR on the genome | 0.59194918 |

Relative frequency (%) of SSR types, by number of repeats, in the Ensete glaucum genome. The graph (Fig. 7A) is based on a total of N = 123884 SSRs detected in the Ensete glaucum genome.

| Repeats % count | 4 | 5 | 6 | 7 | 8 | 9 | 10 | 11 | 12 |
|---|---|---|---|---|---|---|---|---|---|
| Mono | 0 | 0 | 0 | 0 | 0 | 0 | 0 | 0 | 8.13907 |
| Di | 0 | 0 | 0 | 0 | 5.504342772 | 4.811759388 | 4.13209 | 3.37251 | 2.7219 |
| Tri | 0 | 9.516160279 | 4.69229279 | 2.8615479 | 1.861418747 | 1.296374027 | 0.82819 | 0.5263 | 0.36486 |
| Tetra | 0 | 1.452972135 | 0.668367182 | 0.237318782 | 0.112201737 | 0.046817991 | 0.02583 | 0.0226 | 0.00726 |
| Penta | 2.319104969 | 0.781376126 | 0.272835879 | 0.092021569 | 0.048432405 | 0.025830616 | 0.01049 | 0.01292 | 0.00888 |
| Hexa | 0.782183333 | 0.220367441 | 0.065383746 | 0.017758548 | 0.005650447 | 0.001614413 | 0.00323 | 0.00081 | 0 |

| Repeats % count CONTINUED | 13 | 14 | 15 | 16 | 17 | 18 | 19 | 20 | >20 | Total |
|---|---|---|---|---|---|---|---|---|---|---|
| Mono | 6.063737044 | 4.609957702 | 3.541215976 | 2.770333538 | 1.884020535 | 1.191437151 | 0.67886 | 0.41894 | 0.74909 | 30.0467 |
| Di | 2.279551839 | 1.980885344 | 1.708856672 | 1.50140454 | 1.244712796 | 1.100222789 | 0.98318 | 0.83788 | 6.61667 | 38.796 |
| Tri | 0.338219625 | 0.25023409 | 0.174356656 | 0.186464757 | 0.152562074 | 0.157405315 | 0.10897 | 0.10171 | 0.43347 | 23.8505 |
| Tetra | 0.008072067 | 0.005650447 | 0.003228827 | 0.003228827 | 0.003228827 | 0.000807207 | 0.00161 | 0.00081 | 0.00726 | 2.60728 |
| Penta | 0.00484324 | 0.00242162 | 0.003228827 | 0.004036034 | 0.000807207 | 0.00242162 | 0.00323 | 0.00081 | 0.00484 | 3.59853 |
| Hexa | 0.000807207 | 0 | 0.000807207 | 0.001614413 | 0 | 0 | 0 | 0 | 0.00081 | 1.10103 |

TRUNCATED IN PDF Version
